# Histidine exchange sustains LAT1 activity and proliferation in glutamine-addicted breast cancers

**DOI:** 10.64898/2026.04.15.716193

**Authors:** Ersa Gjelaj, Paul C. Driscoll, Asad Mahmood, Josep Tarrago-Celada, Michael Kossifos, Saidu Sesay, James I. MacRae, Mariia Yuneva

## Abstract

L-type amino acid transporter (LAT1) drives the uptake of essential amino acids (EAA) and activation of mTORC1 signaling in cancer cells. Current models propose that glutamine exchange is required for LAT1-dependent EAA transport; however, in many tumours, including MYC-driven cancers, glutamine is simultaneously consumed for bioenergetic and biosynthetic processes. How LAT1 activity is maintained under these conditions remains unclear. Here, we identify histidine as an efficient bidirectional LAT1 substrate that is preferentially utilised under glutamine-limitation in Gln-dependent tumour cells. Histidine uptake is modulated by glutamine availability, revealing an unexpected role for histidine in maintaining amino acid homeostasis. We demonstrate that histidine availability supports LAT1-mediated transport, promotes mTORC1/4E-BP1 signaling, and enhances protein synthesis and supports tumour cell proliferation. Under histidine limitation, MYC and ATF4 induce amino acid transporter expression, including LAT1, to preserve intracellular EAA levels. Importantly, histidine restriction sensitizes tumour cells to LAT1 inhibition, enhancing sensitivity to LAT1 inhibition and reducing tumour burden. Together, our findings establish histidine as a key regulator of LAT1 function and mTORC1 activity, suggesting a potential metabolic vulnerability in glutamine-dependent tumours.

## Introduction

Cancer cells inhabit nutrient-poor, spatially heterogeneous microenvironments and must develop mechanisms to sustain amino acid supply and growth-signaling pathways under chronic metabolic stress. Among these mechanisms, solute carrier (SLC) amino acid transporters play a central role in regulating intracellular amino acid availability and in shaping the activity of nutrient-sensing pathways such as mTORC1. mTORC1 integrates extracellular cues with intracellular nutrient levels to coordinate anabolic growth with nutrient sufficiency, and its activation is tightly coupled to amino acid transport. Thus, the availability of amino acids defines a metabolic bottleneck that many tumours must overcome.

The prevailing model proposed that mTORC1 activation depends on a two-step amino acid transport mechanism involving (i) glutamine import through the high-affinity transporter ASCT2 (encoded by *SLC1A5*) and (ii) glutamine efflux through the LAT1/CD98 (encoded by *SLC7A5*/*SLC3A2*) antiporter, which simultaneously imports extracellular branched- and aromatic-chain EAA, such as leucine^1^. Therefore, intracellular glutamine serves as the efflux substrate that enables EAAs to enter the cell and activate mTORC1, whereas glutamine depletion can limit LAT1-mediated EAA uptake, suppresses mTORC1, and induces autophagy^1^. However, several unresolved issues challenge the universality of this glutamine-centric model. First, LAT1 has relatively low affinity for glutamine, requiring high intracellular concentrations to sustain efficient exchange^2^. Second, glutamine is catabolized for biosynthesis and anaplerosis, particularly in glutamine-addicted tumours^3^. Competition between these biosynthetic demands and LAT1-mediated exchange is therefore expected, particularly in cancer cells with limited capacity for glutamine synthesis. Third, physiological glutamine levels are much lower and variable than those in standard cell culture media, where glutamine is typically supplied at several-fold higher levels than other amino acids^4,5^. This discrepancy raises the possibility that glutamine functions as an exchange substrate *in vitro* but may not reach sufficient levels to do so *in vivo.* Furthermore, ASCT2 knockout does not consistently impair LAT1 activity or mTORC1 signaling in many cancer cell lines, suggesting glutamine is not obligately required for LAT1-mediated EAA uptake^6–8^. Together, these considerations raise the question of how glutamine-dependent cancer cells maintain LAT1 activity, EAA uptake and mTORC1 signaling when glutamine becomes limiting in vivo.

One of the few LAT1 substrates capable of efficient bidirectional transport is the EAA histidine^9^. Histidine also has an exceptionally high affinity towards LAT1, which together with bidirectional transport raise a possibility that under conditions where glutamine availability is constrained, histidine may contribute to sustaining LAT1-mediated amino acid uptake and downstream signaling^2,10^. In this study, we examine amino acid transport in glutamine-dependent cancer cells to evaluate the role of histidine and determine how LAT1 activity is sustained when glutamine availability is limited. We find that histidine is not only abundant and not catabolized in these cells, but also functions as a preferred bidirectional LAT1 substrate, enabling continued uptake of EAAs. This histidine-LAT1 axis supports rapid mTORC1/4E-BP1 activation and translation re-engagement, and its disruption exposes a metabolic vulnerability that may be therapeutically exploited through dietary histidine restriction.

## Results

### MMTV-MYC tumours and tumour cells depend on glutamine catabolism

We first sought to define a tumour context in which glutamine is both heavily utilised and potentially limiting to determine how LAT1-mediated exchange is supported under these conditions. MYC expression has been widely associated with increased glutamine uptake and catabolism, and MYC-driven cancer cells display canonical features of “glutamine-addiction”^3,11^. We therefore used the MMTV-*c-Myc* transgenic mammary tumour model to address these questions.

To characterise glutamine metabolism during MYC-driven mammary gland tumourigenesis, we analysed gene expression profiles across a temporal series comprising normal mammary glands (NMG) from 6-week-old wild-type littermates, hyperplastic mammary glands, and end-stage tumours. Bulk RNA sequencing revealed coordinated changes in the expression of enzymes involved in glutamine metabolism and amino acid transport. Expression of glutaminase (GLS), which catalyses the conversion of glutamine (Gln) to glutamate (Glu), progressively increased during tumour progression, whereas expression of glutamine synthetase (GS/GLUL), which synthesizes glutamine from glutamate, was negatively correlated with MYC-driven tumour progression (Fig. 1a,b). In parallel, multiple amino acid transporters showed positive correlations with tumour progression, including members of the *Slc7*, *Slc43,* and *Slc38* families, several of which are known MYC transcriptional targets (Fig. 1a,b)^12^. Notably, *Slc7a5* expression increased early in hyperplastic mammary glands and remained elevated throughout tumour progression (Fig. 1a,b). Consequently, expression of *Slc3a2* (CD98), which is required for LAT1 trafficking to the plasma membrane, strongly correlated with both MYC-driven tumour progression and *Slc7a5* expression (Fig. 1a,b). Immunoblot analysis of end-stage MMTV-c-*Myc* tumours compared to normal mammary gland tissue confirmed increased expression of GLS1 and the amino acid transporters LAT1 and ASCT2, together with reduced expression of GS (Fig. 1c), consistent with a shift toward net glutamine uptake and catabolism in MYC-driven tumours. Immunohistochemical analysis of LAT1 revealed marked inter- and intra-tumour heterogeneity in plasma membrane localisation (Fig. 1d). Quantitative scoring of LAT1 membrane staining across individual tumours demonstrated a broad distribution of expression levels; however, the majority of tumour cells (74.91 ± 14.21%, n= 14) exhibited detectable LAT1 localisation at the plasma membrane (Fig. 1d). These data indicate that, despite tumour heterogeneity, LAT1 is predominantly positioned at the cell surface in MYC-driven mammary tumour (MTs) cells, suggesting functional amino acid transport capacity.

**Fig. 1:**
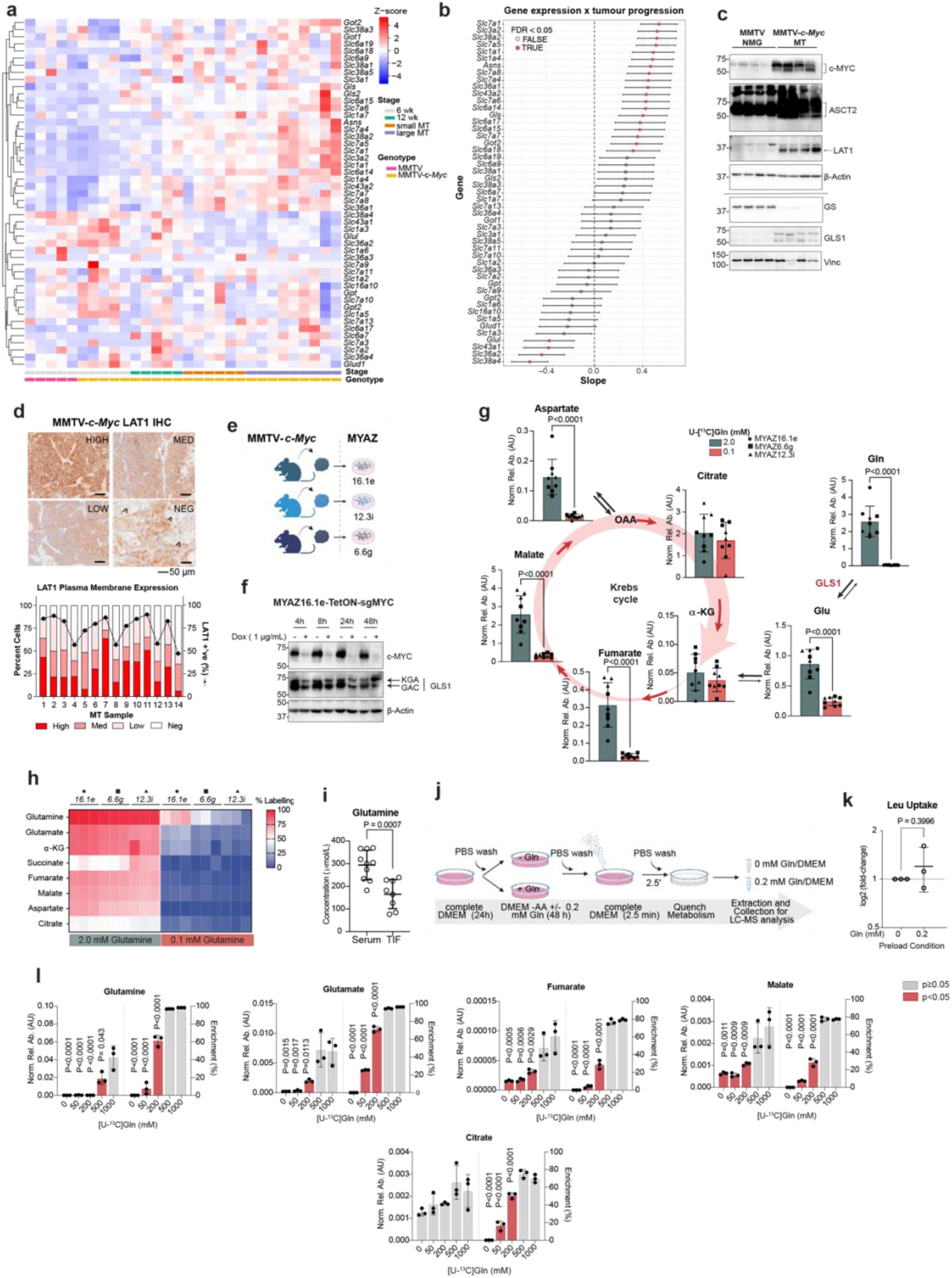
*Slc7a5*/LAT1 expression and glutamine dependence are increased in MYC-driven mammary tumourigenesis. (**a**) Heatmap of *Slc* and glutamine metabolism gene expression from bulk RNA-seq of NMG, hyperplastic mammary glands, and tumours during MMTV-c-*Myc* driven tumourigenesis (n = 5, 5, 5, 6, 9 from left to right). (**b**) Gene-wise association with disease progression. Slopes from gene-wise linear models (expression ∼ stage) with 95% CI. (**c**) WB of indicated proteins in NMG and MMTV-c-*Myc* tumours. (**d**) Representative LAT1 IHC staining (top) and membrane quantification (bottom) in MMTV-c-*Myc* tumours (n = 14). (**e**) Schematic for cell line generation from MMTV-c-*Myc* tumours. (**f**) WB of c-MYC and its transcriptional target Gls1 in MYAZ16.1e clones with Dox-inducible sg*Myc* (n = 2). (**g-h**) LC-MS analysis of cells cultured in 2.0 or 0.1 mM [U^13^C]Gln for 24 h (n = 3 per cell line). (**g**) Relative metabolite abundance. Mean ± s.d.; *p*-values from unpaired two-tailed t-test. (**h**) Heatmap of ^13^C enrichment. (**i**) Absolute Gln concentrations in serums (n = 10) and TIF (n = 8) from mice bearing MYAZ16.1e orthotopic tumours. Mean ± s.d, paired two-tailed t-test. (**j**) A schematic for Gln preloading experiment: cells starved of amino acids (-AA) ± 0.2 mM Gln preload for 48 h, then replenished with DMEM. (**k**) Fold change of Leu levels measured by LC-MS in amino acid-starved cells or those preloaded with 0.2 mM Gln and replenished with DMEM for 2.5 min (n = 3). Mean ± s.d.; unpaired two-tailed t-test. (**l**) Total abundance and ^13^C enrichment of indicated metabolites measured by LC-MS in in amino acid-starved cells or those preloaded with indicated concentrations of [U-^13^C]Gln and replenished with DMEM for 2.5 min (n = 3). Mean ± s.d.; one-way ANOVA with Dunnett’s test for multiple comparisons against 1000 µM. **See also Figure S1.**

To directly examine glutamine utilisation in tumour cells, we established cell lines derived from independent MMTV-c-*Myc* MTs (MYAZ-6.6g, 12.3i, 16.1e; Fig. 1e). To determine whether Gln catabolism in these cells is actively maintained by MYC, we employed a doxycycline-inducible MYC knockout system in MYAZ16.1e cells. Acute loss of MYC results in a partial reduction in GLS expression (Fig. 1f), indicating that MYC activity is necessary to sustain GLS expression. We next traced the fate of Gln carbons using [U-^13^C]Gln. At high Gln availability (2 mM), downstream metabolites including Glu, alpha-ketoglutarate (𝘢-KG), fumarate, malate, and aspartate were primarily maintained by Gln-derived carbon, as indicated by >50% ^13^C enrichment in these Krebs cycle intermediates. (Fig. 1g,h). When MYAZ16.1e tumour cells were cultured at low Gln concentrations (100 µM), both the abundance and labelling of Krebs cycle intermediates were markedly reduced (Fig. 1g,h). Consistent with impaired anaplerosis, tumour cell proliferation significantly decreased at low Gln concentrations, which was rescued by supplementation with dimethyl alpha-ketoglutarate (DMKG) (Fig. S1a,b). Pharmacological inhibition of GLS1 with CB-839 also suppressed proliferation (Fig. S1c). Together, these data establish MMTV-c-*Myc* MTs and derived cell lines as a Gln-addicted model characterised by high LAT1 and ASCT2 expression and a strong reliance on exogenous glutamine to sustain Krebs cycle activity.

Because LAT1-mediated transport requires intracellular exchange substrates, we next asked whether glutamine is available *in vivo* at concentrations sufficient to support Gln-dependent LAT1 activity and leucine uptake. Measurement of Gln concentrations in serum and tumour interstitial fluid (TIF) from orthotopic MYAZ16.1e tumours revealed that circulating Gln levels were 294.3 µM ± 65.4 (n = 10) with Gln concentrations within the TIF being significantly reduced (165.7 µM ± 64.4, n = 8; one outlier removed; Fig. 1i). To model these physiological conditions *in vitro*, MYAZ16.1e cells were subjected to a Gln preloading protocol prior to acute metabolic analysis (Fig. 2j). Preloading cells with 200 µM Gln, corresponding to the concentration measured in the TIF, did not enhance leucine uptake relative to Gln-free conditions (Fig. 2k). Consistent with this observation, [U-^13^C]Gln tracing revealed that intracellular abundance and ^13^C enrichment of multiple Krebs cycle intermediates were markedly reduced at Gln concentrations ≤ 200 µM, whereas glutamine concentrations ≥ 500 µM were required to maintain Krebs cycle metabolite levels and anaplerotic flux (Fig. 2l). Together, these data suggest that Gln concentrations in the tumour microenvironment are lower than those required to sustain intracellular Gln pools sufficient for Krebs cycle anaplerosis and are therefore unlikely to efficiently support glutamine-dependent amino acid exchange for LAT1 substrates such as leucine. These findings raise the possibility that tumour cells with high glutamine demand may rely on alternative amino acids to support LAT1-mediated exchange under Gln-limited conditions. ,

**Fig. 2:**
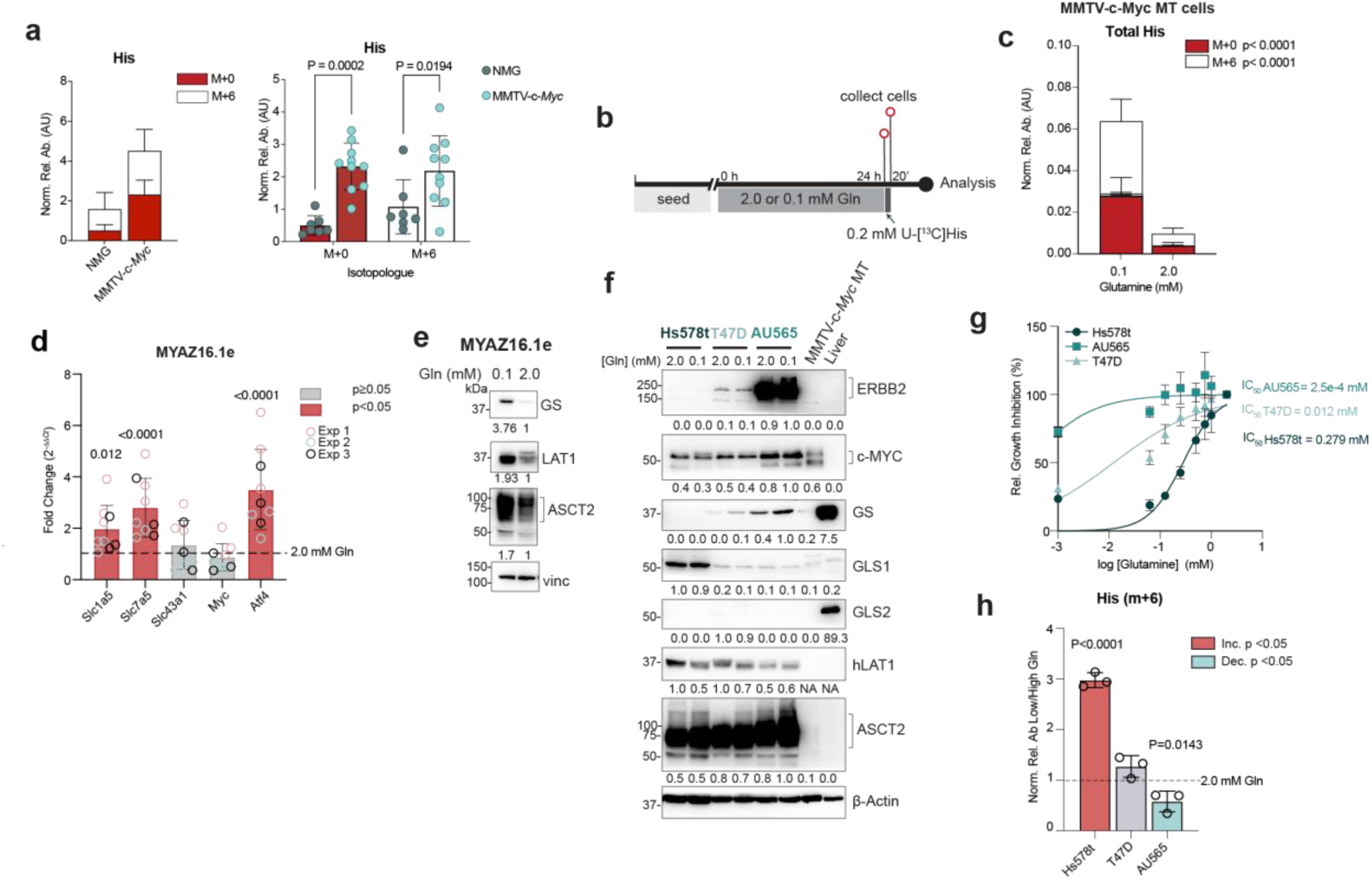
Gln-availability is a determinant of His requirement. (**a)**LC-MS of His levels in NMG (n = 7) and MMTV-c-*Myc* (n = 10) tumours after a 5 min [U^13^C]His bolus. Total abundance (left) and isotopologue distribution (right). Mean ± s.d.; *p*-values from two-way ANOVA (Šídák’s correction). (**b)** Timeline: cells cultured in 2.0 or 0.1 mM Gln (24 h) before 20-min [U^13^C]His administration. (**c)** Total relative His abundance (n = 3). Data are mean ± s.d.; *p*-values from two-way ANOVA (Šídák’s correction). (**d**) RT-qPCR of indicated genes expression in cells cultures in 2.0 mM or 0.1 mM Gln (n = 3), normalized to β-actin and expressed relative to 2.0 mM Gln. Mean ± s.d.; *p*-values from one-way ANOVA (Šídák’s correction). (**e,f**) WB of cells in 2.0 mM or 0.1 mM Gln (24 h), n = 3; in (**f**) MMTV-c-*Myc* tumour and liver tissue are used as controls. (**g**) Relative confluency of cells in 0-2.0 mM Gln (96 h), normalized to 2.0 mM Gln; IC50 values from non-linear fit of log[Gln] vs. normalized response (n = 2). (**h**) LC-MS of cells in 2.0 mM or 0.1 mM Gln (24 h), followed by a 10 min [U^13^C]His pulse (n = 3). Labelled His abundance is relative to 2.0 mM Gln. Mean ± s.d.; *p*-values from one-way ANOVA (Šídák’s correction). **See also Figure S2.**

### Glutamine availability influences histidine uptake in glutamine-dependent tumour cells

LAT1 exhibits its highest affinity towards histidine (His; Km 12 µM)^2,10^ and, unlike other EAAs, is capable of mediating bidirectional His transport^9^. We therefore asked whether His uptake is altered in MYC-driven tumours and tumour cells that exhibit high Gln demand and experience limited Gln availability. To assess His uptake *in vivo*, we administered a short intravenous bolus of uniformly labelled His, [U-^13^C]His, to MMTV-c-*Myc* tumour-bearing mice and quantified histidine abundance in MTs and NMGs from age-matched wild-type controls. Both unlabelled (m+0) and labelled (m+6) His were significantly increased in MYC-driven MTs compared with NMG control (Fig. 2a), indicating enhanced His uptake and accumulation in tumours. While the mechanisms underlying His accumulation *in vivo* remain to be fully defined, these data suggest that MYC-driven tumours exhibit increased His acquisition relative to NMG tissue.

We next examined whether His uptake is influenced by Gln availability in MYC-driven tumour cells. MMTV-c-*Myc* MT (MYAZ-6.6g, 12.3i, 16.1e) cells were cultured under Gln-replete (2.0 mM) or Gln-limited (100 µM) conditions for 24 h prior to acute tracing with [U-^13^C]His at 200 µM, a concentration of His in DMEM (Fig. 2b). Under Gln-limited conditions, tumour cells exhibited significantly increased total His abundance and enhanced incorporation of ^13^C-labelled His compared with cells cultured in Gln-replete medium (Fig. 2c). Gln limitation was accompanied by increased expression of the amino acid transporters *Slc1a5* and *Slc7a5*, induction of the stress- responsive transcription factor *Atf4*, and increased protein abundance of ASCT2 and LAT1 (Fig. 2d,e), consistent with activation of the integrated stress response and adaptive amino acid uptake.

Previous work by Nicklin *et al*., demonstrated that cells capable of synthesizing Gln are less dependent on extracellular Gln to activate mTORC1 in response to EAA availability via LAT1 - facilitated uptake^9^. We therefore asked whether the ability to synthesise Gln influences His uptake under Gln-limited conditions. To address this, we compared breast cancer cell lines with differing Gln metabolic profiles but comparable MYC expression (Fig. S2a and Fig. 2f). Hs578t cells, which express high levels of GLS and lack detectable GS, exhibited strong dependence on extracellular Gln for proliferation (Fig. 2f,g). In contrast, T47D and AU565 cells expressed GS, with AU565 cells showing the highest GS abundance and the least sensitivity to Gln deprivation (Fig. 2f,g). Consistent with these differences, [U^13^C]Gln tracing revealed greater reliance on Gln-derived carbon for Krebs cycle anaplerosis in Hs578t cells, whereas T47D and AU565 cells showed increased contributions from alternative carbon sources (Fig. S2b,c).

We next assessed His uptake in these cell lines under Gln-limited conditions. In Gln-dependent Hs578t cells, [U-^13^C]His uptake was significantly increased under Gln-limited conditions relative to Gln-replete conditions (Fig. 2h). In contrast, His uptake was unchanged in T47D cells and reduced in AU565 cells under Gln limitation (Fig. 2h). Thus, enhanced His uptake correlated with cellular dependence on exogenous Gln and inversely with the capacity for Gln synthesis.

Together, these data indicate that His uptake is selectively increased in tumours and tumour cells where Gln availability is constrained and Gln demand is high. In contrast, tumour cells with reduced reliance on Gln catabolism or increased capacity for Gln synthesis exhibit diminished His uptake under Gln-limited conditions. These findings suggest that His may fulfil a distinct role under conditions of Gln-limitation.

### Histidine demand in MYC-driven mammary tumours is independent of His catabolism

We next examined whether increased His uptake under Gln-limited conditions reflects enhanced His catabolism or instead supports a catabolism-independent mechanism. His degradation can generate glutamate and donate one-carbon units to the folate cycle via formiminotransferase cyclodeaminase (FTCD), histidine catabolism could, in principle, compensate for reduced glutamine availability by sustaining glutamate production (Fig. S3a).

To establish the physiological context of His catabolism, we first analysed publicly available RNA-seq datasets from normal tissues of adult FVB/N mice^13^. Expression of genes encoding His catabolic enzymes, including *Hal*, *Amdhd1*, *Uroc1*, and *Ftcd*, was largely restricted to the liver, with minimal expression detected in other tissues, including the mammary gland (Fig. S3b) . Consistent with this pattern, infusion of [U-^13^C]His into healthy FVB/N mice resulted in high levels of labelled His in serum and across multiple tissues, however, incorporation of His-derived carbon into glutamate was detected only in the liver (Fig. S3c). Labelled glutamine was also observed in the liver, likely reflecting hepatic GS activity, with lower levels detected in peripheral tissues, consistent with systemic release of liver-derived glutamine (Fig. S3c).

We next assessed His catabolism in MYC-driven MTs. In MMTV-*c-Myc* mice, His catabolic enzymes were expressed in the liver but were undetectable in MTs (Fig. S3d). Following infusion of [U-^13^C]His into tumour-bearing mice, both tumours and livers accumulated labelled His, however, His derived formiminoglutamate (FIGLU) and glutamate were detected exclusively in the liver and not in tumours (Fig. S3e-h), indicating an absence of active His catabolism in MYC-driven mammary tumours *in vivo*.

To further exclude His catabolism in tumour cells, we cultured MYAZ16.1e MT cells with [U-^13^C]His under conditions of Gln or methionine limitation. Across all conditions, the majority of intracellular His was labelled, confirming efficient uptake (Fig. S3i). However, Gln limitation resulted in reduced intracellular glutamate levels without detectable incorporation of His-derived carbon into glutamate, indicating that His does not compensate for Gln deprivation through catabolic conversion to glutamate (Fig. S3j). Similarly, under methionine-limited conditions, His did not contribute to methionine regeneration or restore SAM levels or histone methylation, even in the presence of homocysteine and vitamin B_12_ (Fig. S3k-m).

Collectively, these data demonstrate that His catabolism is inactive in MYC-driven MTs and tumour-derived cells and does not account for the increased His uptake observed under Gln-limited conditions. These findings support a catabolism-independent role for His in Gln-dependent tumours and prompted us to investigate whether His instead contributes to amino acid exchange through LAT1.

### His is the preferential LAT1 exchange substrate in MYC-driven cancer cells

Given that MYC-driven tumour cells do not catabolise His, we next asked whether His instead functions as an amino acid exchange substrate to support LAT1-mediated transport under Gln-limited conditions. To address this, we first examined His efflux following acute His loading under Gln-replete (2.0 mM) and Gln-limited (100 µM) conditions. MYAZ16.1e cells were pulsed with [U-^13^C]His for 20 min, washed, and replenished with His-free DMEM for 10 min prior to collection of cells and medium. His loading resulted in a marked increase in intracellular His, which was rapidly depleted following medium replenishment (Fig. 3a). Concomitantly, labelled His accumulated in the extracellular medium (Fig. 3b), indicating active His efflux. Consistent with earlier observations of enhanced His uptake under Gln limitation (Fig. 2), both His uptake and efflux were significantly increased in cells cultured in low Gln compared with Gln-replete conditions (Fig. 3a,b). These results indicate that His undergoes rapid bidirectional transport in MYC-driven tumour cells, which is enhanced under Gln-limited conditions.

**Fig. 3:**
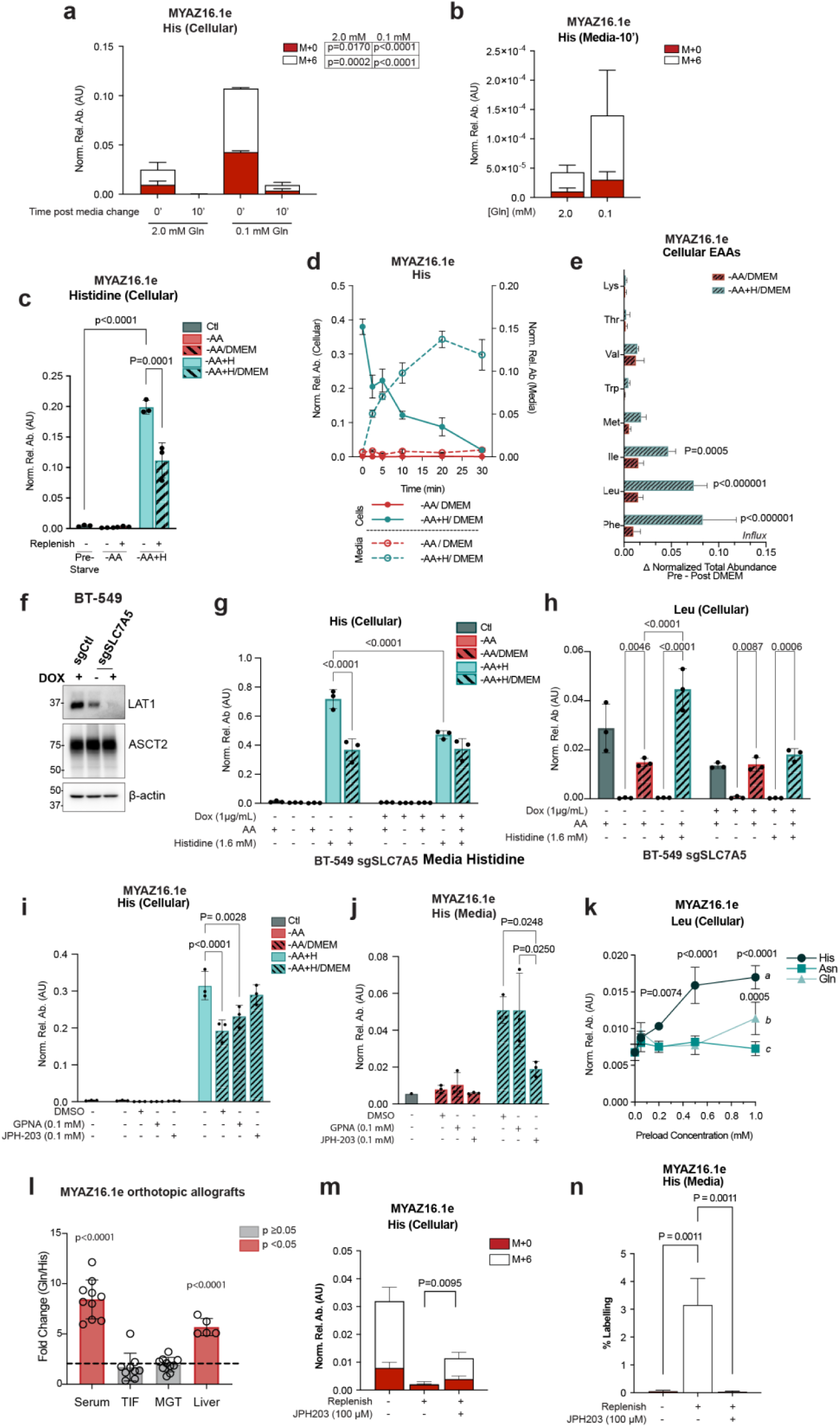
His is an efficient LAT1 exchange factor. (**a,b**) Total relative abundance of (**a**) cellular and (**b**) media His in cells cultured in 2.0 or 0.1 mM Gln for 20 min and replenished with media for 10 min (n = 3). Mean ± s.d.; *p*-values from two-way ANOVA (Šídák’s correction). (**c-e**) Cells starved of amino acids (-AA) ± 1.6 mM His preload (-AA+H) for 48 h, then replenished with DMEM (n = 3): (**c**) Cellular His relative abundance; (**d**) Time-course of relative His abundance in cells and media; (**e**) Intracellular EAA influx, calculated as the difference in abundance post- and pre- replenishment. (**f**) WB of LAT1 in BT-549 sgSLC7A5 ± Dox (48 h, 1 µg/ml, n = 3). (**g**, **h**) LC-MS of His-preloaded BT-549 sgSLC7A5 cells (48 h, 1.6 mM), replenished with DMEM (2.5 min; n = 3): (**g**) His abundance; (**h**) Leu abundance. (**i**, **j**) Relative abundance in cells preloaded with 1.6 mM His, then replenished ± inhibitors (2.5 min, n = 3): (**i**) Cellular His; (**j**) Media His. (**c**-**j**) Mean ± s.d.; *p*-values from two-way ANOVA (Šídák’s correction). (**k**) Leu uptake in cells preloaded with His, Asn, or Gln (48 h), replenished with DMEM (2.5 min, n = 3). Mean ± s.d.; *p*-values from two-way ANOVA (Šídák’s correction). Different letters indicate significant differences: *a* vs. *b* (p<0.0001), *a* vs. *c* (p<0.0001), *b* vs. *c* (p = 0.0014). (**l**) Gln/His fold difference for absolute His and Gln concentrations in MYAZ16.1e orthotopic tumours (n = 10), liver tissue (n = 4), serum (n = 10), and TIF (n = 9). (**m,n**) Relative abundance in cells preloaded with 0.05 mM [U^13^C]His and 0.3 mM [U^13^C]Gln (48 h), replenished ± JPH203 for 2.5 min (n =3): (**m**) Cellular His; (**n**) Labelled His in media. Mean ± s.d.; *p*-values from one-way ANOVA (Šídák’s correction). **See also Figure S3 and S4.**

To determine whether His efflux supports the uptake of other amino acids, we established a His preloading paradigm. MYAZ16.1e cells were amino acid starved (-AA) for 48 h in the presence or absence of 1.6 mM His (-AA+H), allowing intracellular His pools to reach steady state, before replenishment with His-free DMEM for 2.5 minutes. Following His preloading, intracellular His levels were markedly elevated and rapidly decreased upon medium replenishment, consistent with His efflux (Fig. 3c). Time-course analysis confirmed rapid His depletion from cells and accumulation in the medium within 30 minutes of replenishment (Fig. 3d). We next quantified uptake of other amino acids 2.5 min after replenishment. His-preloaded cells exhibited increased influx of multiple EAAs, particularly leucine, phenylalanine, and isoleucine (Fig. 3e). In contrast, uptake of serine, glycine, threonine and valine were unchanged relative to amino acid starved cells (Fig. S4a). Notably, there was significantly increased glutamine uptake in amino-acid starved cells compared to His-preloaded cells (Fig. S4a).

To directly test whether His-mediated amino acid exchange requires LAT1, we disrupted *SLC7A5* using a doxycycline-inducible CRISPR-Cas9 system in BT-549 cells, a MYC-amplified triple-negative breast cancer cell line characterised by high *GLS1* and low *GLUL* expression (Fig. S2a, Fig. 3f). In control (Dox-off) cells, His preloading followed by medium replenishment resulted in significant intracellular His depletion (Fig. 3g). In contrast, LAT1-deficient (Dox-on) cells displayed reduced intracellular His accumulation following preloading and failed to further deplete His upon replenishment (Fig. 3g). Similarly, Leu uptake following His preloading was observed in control cells but was abolished in LAT1-deficient cells (Fig. 3h). Comparable effects were observed for other high-affinity LAT1 substrates, including tryptophan, isoleucine, methionine, phenylalanine, and tyrosine (Fig. S4b). Consistent with these genetic perturbations, pharmacological inhibition of LAT1 with JPH203 in MYAZ16.1e cells blocked His efflux following medium replenishment, whereas inhibition of ASCT2 with GPNA had no effect (Fig. 3i,j). Moreover, the enhanced uptake of EAAs and tyrosine observed in His-preloaded cells was strongly suppressed by JPH203 (Fig. S4c). Together, these data indicate that LAT1 is required for His efflux and for His-dependent uptake of EAA. Incomplete inhibition of His uptake upon inhibiting LAT1 expression is also consistent with a model in which His uptake and His-mediated exchange are mediated by distinct transport systems, with LAT1 functioning primarily as the efflux and exchange component.

We next compared His with previously described amino acid exchange substrates of LAT1, Gln and asparagine (Asn)^9,12^. MYAZ16.1e cells were amino acid starved and preloaded with His, Asn, or Gln across a range of concentrations (50 µM - 1 mM), followed by replenishment with His, Asn, or Gln-free DMEM, respectively, for 2.5 min. His preloading enhanced leucine uptake in a concentration-dependent manner, with a significant effect observed at 200 µM (Fig. 3k). In contrast, Asn preloading failed to enhance leucine uptake even at supraphysiological concentrations tested (i.e., 1 mM), which is well above reported human plasma/blood Asn concentrations of approximately 50-60 µM^14^, whereas Gln preloading promoted leucine uptake only at high concentrations and less efficiently than His (Fig. 3k). Similar patterns were observed for other LAT1 substrates (Fig. S4d), whereas threonine and serine were preferentially increased following Gln or Asn preloading (Fig. S4d). To further define LAT1 dependence, we assessed amino acid uptake in preloaded cells following LAT1 inhibition in cells preloaded with 1 mM of His, Gln, or Asn or in the absence of all amino acids. In amino acid starved cells, JPH203 had minimal effects on amino acid uptake. In contrast, His-preloaded cells exhibited pronounced LAT1-dependent uptake of EAAs, which was strongly inhibited by JPH203 (Fig. S4e). Gln-preload cells facilitated LAT1-mediated exchange of some substrates, but less efficiently than His, as indicated by lower LAT1 substrate levels in replenished Gln-preloaded compared to His-preloaded cells (Fig. S4e). In contrast, amino acid uptake in Asn-preloaded cells were largely LAT1-independent (Fig. S4e). These results indicate that, in Gln-dependent tumour cells, Gln availability for LAT1 mediated exchange is constrained by its low affinity for LAT1 and its rapid metabolic utilisation, such that Gln can function as an effective exchange substrate only when present at supraphysiological concentrations.

We next confirmed the relationship between Gln and His as LAT1 exchange factors for physiological concentrations of these amino acids. Gln concentrations in the serum of healthy adults (500 to 800 µM)^15^ are higher than those of His (70 to 120 µM)^16^. Moreover, amino acid availability in the serum may differ from that in the TME. We compared Gln levels in the serum and TIF of mice with orthotopic MYAZ16.1e tumours (Fig. 1i), as well as in tumours and livers, to His levels in the serum, TIF, tumours, and livers of these animals (Fig. 3l). As expected, serum Gln levels were significantly higher than His levels (Fig. 3l). However, in the TIF, there was no significant difference between His and Gln concentrations (Fig. 3l). Notably, His levels were significantly higher in the TIF (125.4 µM ± 39.6, *n*=9) than in the serum, whereas Gln concentration in the TIF (165.7 µM ± 64.4, n=8) were significantly lower than in serum (Fig. S4f and Fig. 1i). Furthermore, while Gln and His concentrations in MTs were comparable, Gln levels were significantly higher in the liver tissues (Fig. 3l). These findings suggest that MYC-driven MTs exhibit an increased demand for Gln catabolism and exist in a Gln-limited environment. Consequently, His which is present at higher concentrations in the tumour environment and within MTs can serve as an effective LAT1 exchange substrate under these conditions.

Next, to directly compare His and Gln as LAT1 exchange substrates under identified physiologically relevant conditions, we performed competitive preload experiments using MYAZ16.1e tumour cells. Cells were preloaded with both [U-^13^C]Gln (300 µM) and [U-^13^C]His (50 µM) under amino acid-free conditions for 48 h, followed by replenishment with DMEM ± JPH203 for 2.5 min. Consistent with previous observations, intracellular His levels declined upon DMEM replenishment, with a concomitant increase in labelled His in the media (Fig. 3m,n). However, in the presence of JPH203, His efflux was blocked and labelled His was no longer detected in the media (Fig. 3m,n, Fig. S4g). In contrast, intracellular Gln levels remained largely unchanged upon DMEM replenishment, and LAT1 inhibition did not affect Gln depletion, as indicated by the absence of labelled Gln in the media (Fig. S4h-i). These results demonstrated that even at lower concentrations, His showed more robust LAT1-dependent exchange than Gln under these experimental conditions. Given that Gln levels in the TIF and MTs are not significantly higher than those of His (Fig. 3l), this underscores the potential for His to play a prominent role in sustaining LAT1-mediated amino acid uptake in glutamine-dependent MTs, such MYC-driven MTs.

### His availability primes tumour cells for increased mTORC1/4E-BP1 signaling

LAT1 sustains intracellular pools of EAAs, including leucine, which regulates mTORC1 activity through cytoplasmic sensors such as SESTRIN2 and SAR1B^17,18^. Given our observation that His availability enhances EAA uptake, we next examined whether His availability influences mTORC1 activation following amino acid replenishment. BT-549 cells were starved of amino acids for 48 h in the presence or absence of His and then replenished with complete DMEM. mTORC1 activation was assessed by monitoring phosphorylation of RPS6 and 4E-BP1. His-preloaded cells exhibited enhanced phosphorylation of 4E-BP1 relative to controls, consistent with more rapid re-engagement of mTORC1 signaling upon replenishment (Fig. 4a). In contrast, phosphorylation of RPS6 was reduced in His-preloaded cells compared to controls (Fig. 4a), suggesting that His-facilitated amino acid uptake differentially influences mTORC1 downstream outputs, potentially reflecting differences in signaling kinetics or pathway sensitivity.

**Fig. 4:**
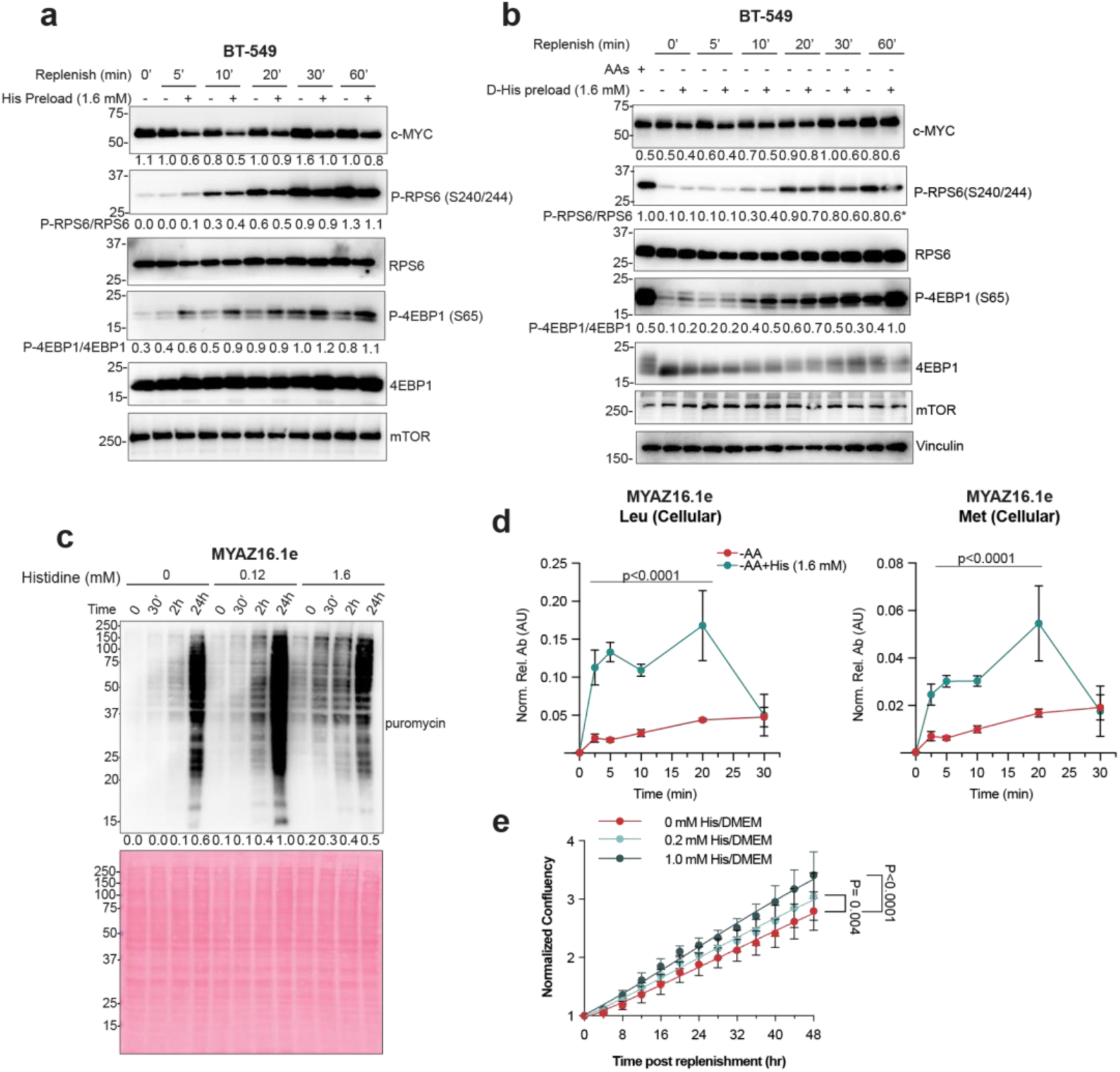
His availability enhances mTORC1/4E-BP1 activation via LAT1-mediated EAA exchange. (**a,b**) WB of replenished cells after amino acid starvation with (**a**) 1.6 mM His ( n = 3) or (**b**) 1.6 mM D-His (n = 2) for 48 h. Protein levels are normalized to mTOR/Vinc and expressed relative to the maximum. (**c**) WB of puromycin incorporation in cells starved or preloaded with 0.12 mM or 1.6 mM His (48 h), then replenished with DMEM. Protein levels normalized to Ponceau and expressed relative to the maximum. A representative of 2 independent replicates. (**b)** Time-course of relative abundance of intracellular Leu (left) and Met (right) in DMEM-replenished cells after His preloading (1.6 mM, 48 h). 0 min represents pre-replenishment levels. Mean ± s.d.; *p*-value from two-way ANOVA (Šídák’’s correction). (**e**) Confluency of DMEM-replenished cells preloaded with His, normalized to pre-replenishment confluency (n = 3). Gompertz growth curve applied. Mean ± s.d.; *p-*values from two -way ANOVA (Dunnett correction). **See also Figure S5.**

To determine whether mTORC1 activation depends on His-mediated EAA uptake via LAT1, we next assessed mTORC1 activation in cells preloaded with D-His. LAT1 efficiently transports L-His but exhibits limited stereoselectivity for D-His^2^. Although D-His accumulated intracellularly during starvation, it was not efficiently effluxed following DMEM replenishment, and uptake of LAT1 substrates including leucine, isolecuine, phenylalanine, and tyrosine was not increased relative to amino acid starved controls (Fig. S5a-c). Consistent with the absence of LAT1-mediated exchange, D-His preloading failed to enhance phosphorylation of either 4E-BP1 or RPS6 following replenishment (Fig. 4b). These data indicate that His-dependent activation of mTORC1 signaling correlates with LAT1-mediated essential amino acid uptake.

Given that 4E-BP1 phosphorylation promotes translation initiation, we next examined the impact of His availability on protein synthesis. Using the SUnSET assay, nascent protein synthesis was measured in MYAZ16.1e cells preloaded with increasing concentrations prior to replenishment with complete DMEM. His-preloaded cells initiated protein synthesis more rapidly than amino acid starved controls upon amino acid replenishment, with increased puromycin incorporation evident as early as 30 minutes after replenishment and scaling His concentration (Fig. 4c). Consistent with heightened translational activity, intracellular levels of EAAs, including leucine and methionine, declined rapidly following replenishment in His-preloaded cells (Fig. 4d), consistent with increased utilisation. Finally, we assessed whether His availability influences proliferative capacity. Preloading cells with His resulted in a concentration-dependent increase in rate of proliferation (Fig. 4e). Although His concentrations used in these preloading experiments exceed those typically measured *in vivo*, these findings indicate that elevated intracellular His can enhance LAT1-dependent amino acid uptake and accelerate nutrient-responsive activation of mTORC1/4E-BP1 signaling and protein synthesis. Together, these data suggest that His availability can prime tumour cells for rapid recovery of amino acid-responsive signaling and translational activity following nutrient depletion, a property that may be advantageous in Gln- limited environments where efficient amino acid exchange through LAT1 becomes critical.

### Histidine limitation increases the expression of amino acid transporters to maintain amino acid homeostasis

Inhibition of LAT1 is known to activate the integrated stress response (ISR), underscoring its central role in amino acid homeostasis^7,19^. Given that His functions as an efficient LAT1 exchange substrate in Gln-dependent tumour cells, we next asked whether His uptake represents a metabolic vulnerability in this context. To address this, we examined the effects of His limitation on mTORC1 signaling and ISR activation. Reducing extracellular His resulted in a concentration-dependent attenuation of mTORC1 activity, as indicated by decreased RPS6 phosphorylation, together with robust induction of ATF4, consistent with activation of the ISR (Fig. 5a). Notably, despite reduced mTORC1 signaling and ATF4 induction at low His concentrations (50 µM), bulk protein synthesis, assessed by puromycin incorporation, remained largely preserved (Fig. 5a), suggesting that the early responses to His limitation are not driven solely by impaired translational capacity. His limitation also elicited a time-dependent suppression of mTORC1 signaling accompanied by sustained ATF4 induction (Fig. 5b). During this period, MYC protein levels increased (Fig. 5b), suggesting a regulatory link between His availability and MYC expression. Consistent with ISR activation, His deprivation induced expression of the amino acid transporter ASCT2, a known downstream target of the GCN2/eIF2𝘢/ATF4 pathway^20^. Both glycosylated (Glyc) and total ASCT2 protein levels increased under His limitation and correlated with induction of ATF4 and MYC expression (Fig. 5c and Fig. S6a).

**Fig. 5:**
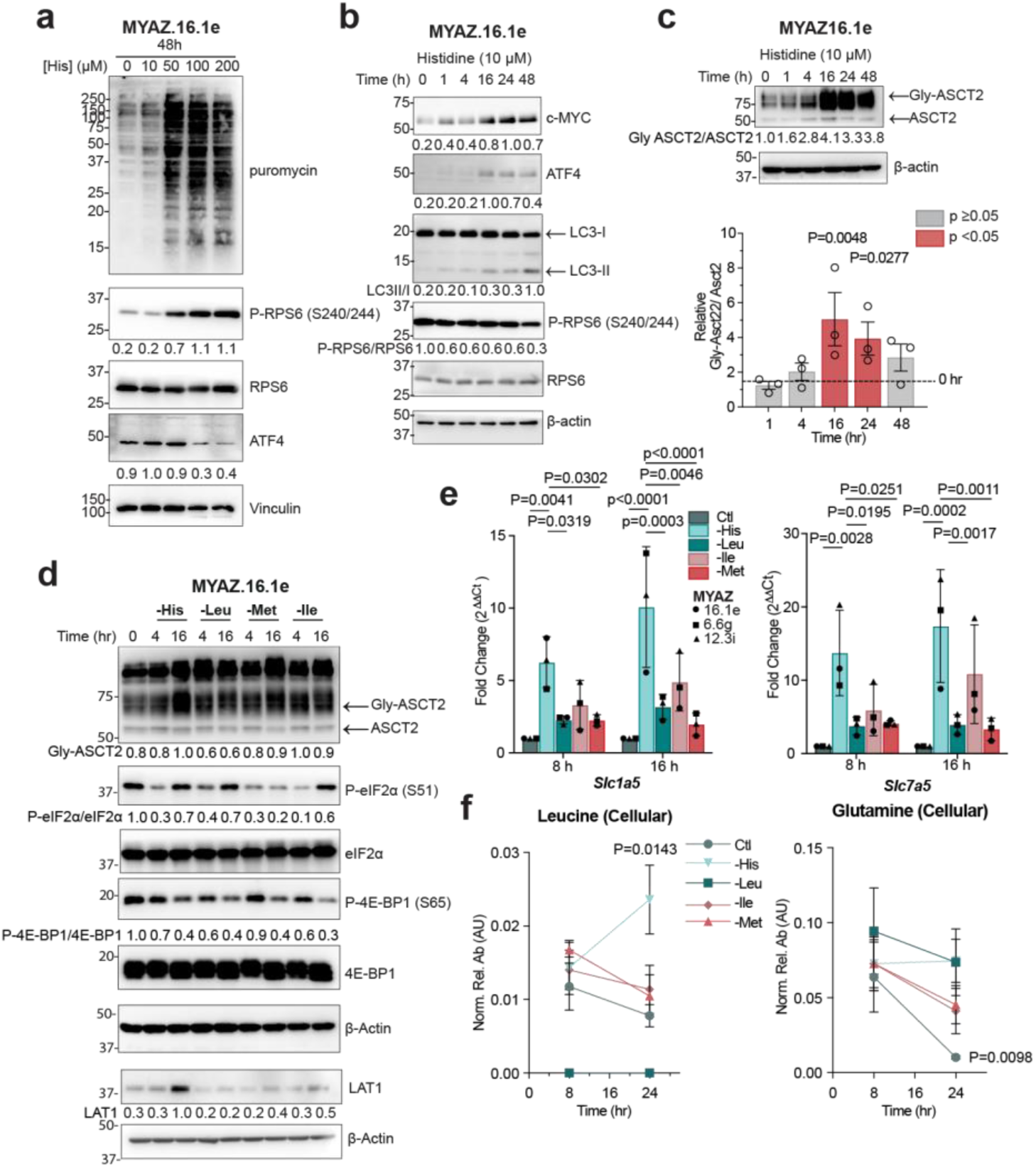
His limitation elicits a stronger amino acid response than other EAAs. (**a, b**) WB of cells cultured in (**a**) 0 to 200 µM His for (48 h, n = 3) or (**b**) 10 µM His (up to 48 h, n = 3). Protein levels normalized to Vinculin or β-actin, expressed relative to the maximum. (**c**) WB (top) and quantification (bottom) of (Glyc) and un-glycosylated ASCT2 in cells cultured in 10 µM His (up to 48 h, n = 3). Protein levels normalized to β-actin, expressed relative to His-replete (0 h). Mean ± s.d.; *p*-values from one-way ANOVA (Šídák’s correction). (**d**) WB of cells cultured in DMEM depleted of His, Leu, Met, or Ile (n = 2). Protein levels normalized to β-actin and expressed relative to the maximum. (**e**) RT-qPCR of cells cultured in complete (Ctl), or His-, Leu-, Ile-, or Met-free DMEM (n = 3), normalized to β-actin and expressed as fold change relative Ctl. Data are mean ± s.d.; *p*-values from two-way ANOVA (Dunnett’s correction). (**f**) Relative abundance of Leu (left) and Gln (right) in cells cultured in complete (Ctl), or His-, Leu-, Ile-, or Met-free DMEM (n = 3), comparing 8 h to 24 h. Mean ± s.d.; *p*-values from two-way ANOVA (Šídák’s correction). **See also Figure S6.**

Because deprivation of several EAAs can activate the ISR^21^, we next compared the effects of His deprivation to that of other high-affinity LAT1 substrates, leucine, isoleucine, and methionine. MYC-driven tumour cells were deprived of His, Leu, Ile, or Met for 4 or 16 h, followed by analysis of mTORC1 and ISR signaling. Deprivation of each amino acid attenuated mTORC1 activity, as reflected by reduced phosphorylation of 4E-BP1, and/or activated the ISR, as indicated by induction of eIF2𝘢 phosphorylation (Fig. 5d). Notably, His deprivation elicited a more pronounced induction of Glyc-ASCT2 than deprivation of Leu, Ile, or Met, which was accompanied by increased localisation of ASCT2 at the plasma membrane (Fig. 5d and Fig. S6b). His deprivation also resulted in stronger induction of LAT1 protein compared with deprivation of Leu, Ile, or Met (Fig. 5d). To determine whether these effects were transcriptionally regulated, we assessed expression of *Slc1a5* and *Slc7a5* following deprivation of His, Leu, Ile, or Met. His deprivation induced *Slc1a5* and *Slc7a5* to a greater extent than deprivation of the other amino acids at both 8 h and 16 h (Fig. 5e), indicating that His availability disproportionately influences transporter expression relative to other LAT1 substrates.

Finally, we examined intracellular amino acid profiles under His limitation. Despite suppression of mTORC1 signaling, His-deprived cells maintained or increased intracellular levels of multiple LAT1 substrates, including leucine, isoleucine, phenylalanine, tryptophan, and tyrosine (Fig. 5f and Fig. S6c). In addition, levels of arginine and lysine, which are taken up independently of LAT1 were also increased during His deprivation compared with deprivation of other EAAs (Fig. S6c). Notably, intracellular Gln levels were maintained during His deprivation, whereas Gln levels declined following deprivation of isoleucine or methionine, or in amino acid-replete control conditions (Fig. 5f).

Together, these data suggest that His limitation perturbs global amino acid homeostasis, triggering a compensatory induction of amino acid transporters that preserves intracellular amino acid pools while simultaneously activating stress signaling pathways. This distinct response to His deprivation supports the notion that His availability represents a specific metabolic constraint in Gln-dependent tumour cells.

### MYC and ATF4 mediate the expression of amino acid transporters upon histidine deprivation

The coordinated induction of MYC and ATF4 under His-limited conditions prompted us to examine their respective contributions to the transcriptional response to His deprivation. ATF4 and MYC are known to co-occupy regulatory regions of numerous metabolic genes, including multiple members of the SLC transporter family, with their relative contributions varying across cellular contexts^12,22,23^. To determine whether His limitation elicits a similar transcriptional program in cells with low basal MYC expression, we used 67NR cells, a murine MT cell line characterized by low endogenous MYC levels.

67NR cells were deprived of His, Ile, Leu, or Met for 16 h, and transcript expression of c*-Myc*, *Slc1a5*, *Slc7a5*, and *Slc38a1* was assessed. His deprivation selectively induced transcription of c-*Myc*, *Slc1a5*, and *Slc7a5* compared with amino acid-replete controls and with deprivation of other EAAs (Fig. 6a), mirroring the response observed in MYC-driven MT cells. Consistent with this, His limitation increased MYC, ASCT2, and LAT1 protein abundance in 67NR cells (Fig. 6b). Induction of these transporters occurred in a His concentration-dependent manner and was accompanied by ATF4 induction (Fig. 6c). Notably, pharmacological inhibition of transcription or translation with Actinomycin D (ActD) or cycloheximide (CHX) abolished MYC induction under His limitation (Fig. 6d), indicating that increased MYC protein levels reflect transcriptional regulation rather than altered protein stability.

**Fig. 6:**
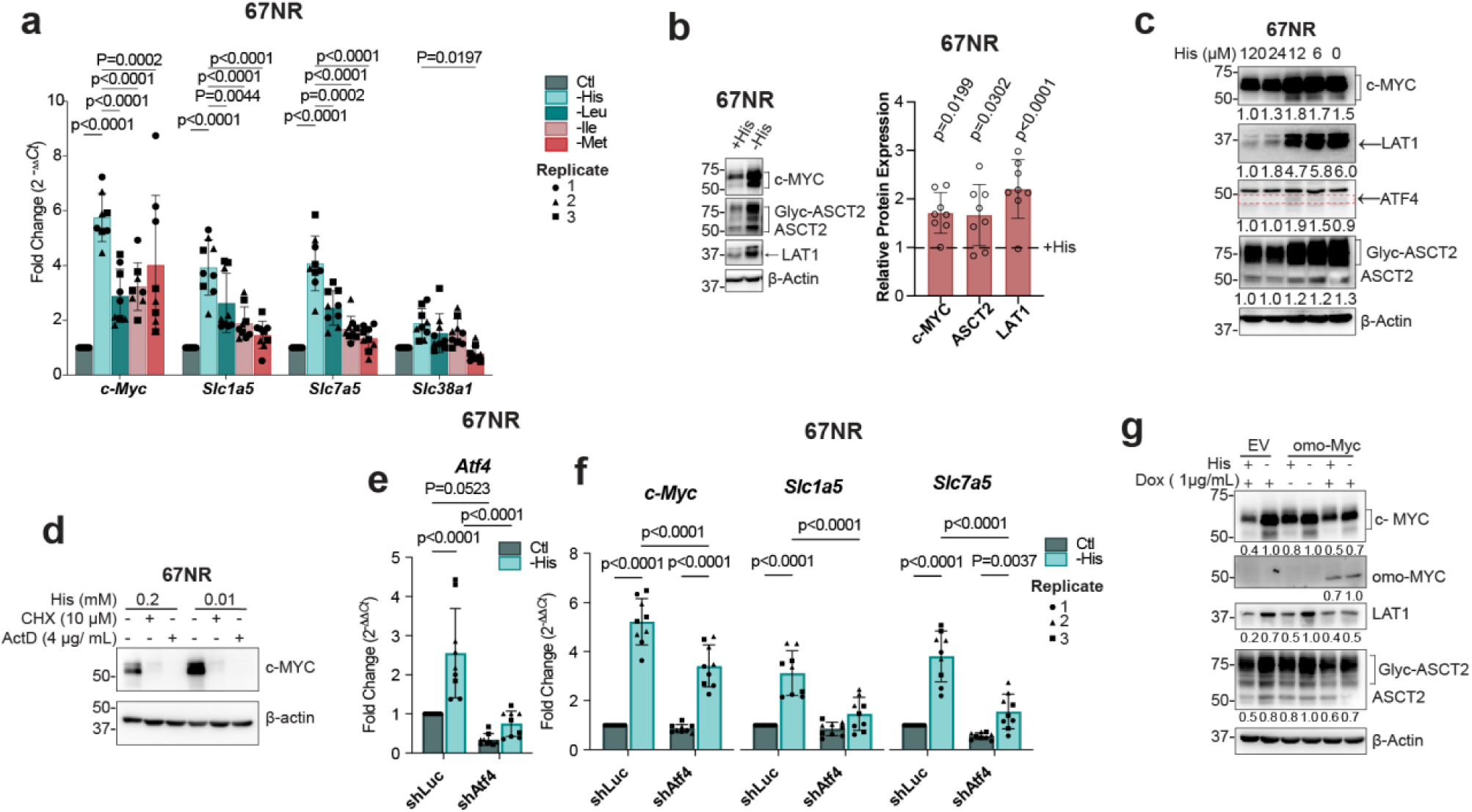
His limitation amplifies MYC- and ATF4-dependent transcriptional responses. (**a**) RT-qPCR of 67NR cells in complete (Ctl), or His-, Leu-, Ile-, or Met-free DMEM (n = 3), normalized to β-actin and expressed as fold change relative Ctl. Mean ± s.d.; *p*-values from two-way ANOVA (Dunnett’s correction). (**b**) WB (left) and quantification (right) of protein expression in cells ± His (24 h, n = 8), normalized to β-actin. Mean ± s.d.; *p*-values from one-way ANOVA (Šídák’s correction). (**c**) WB of cells cultured in 0-120 µM His (24 h, n = 3). Protein levels normalized to β-actin and expressed relative to the maximum. (**d**) WB of cells cultured in 0.2 mM or 0.01 mM His ± CHX or ActD (24 h, n = 3). (**e,f**) RT-qPCR of (**e**) *Atf4* knockdown efficiency and (**f**) gene expression in shLuc or shAtf4 cells ± His (16 h), normalized to β-actin and expressed relative to shLuc + His. Mean ± s.d.; *p*-values from two-way ANOVA (Šídák’s correction). (**g**) WB of cells with Dox-inducible omo-MYC or EV control, ± His or Dox (24 h, n = 1).

Given the strong induction of ATF4 under His limitation, we next assessed its contribution to the transcriptional response. Knockdown of ATF4 in 67NR cells reduced basal *Atf4* transcript levels and blunted its induction upon His deprivation (Fig. 6e). Although c*-Myc*, *Slc1a5*, and *Slc7a5* transcripts were still induced in ATF4-depleted cells, the magnitude of induction was significantly reduced compared with shLuc controls (Fig. 6f), indicating that ATF4 contributes to, but is not solely responsible for, transcriptional upregulation of amino acid transporters under His limitation.

We next evaluated the contribution of MYC activity to this response. To inhibit endogenous MYC function, we generated 67NR cells expressing dox-inducible omo-MYC, a dominant-negative MYC variant that interferes with MYC DNA binding and transcriptional activity. In empty-vector cells and in omo-MYC expressing cells cultured without doxycycline, His deprivation induced MYC, LAT1, and ASCT2 protein expression, consistent with the response observed in the parental cells (Fig. 6g). In contrast, induction of omo-MYC attenuated His-dependent upregulation of MYC and LAT1 (Fig. 6g). These data indicate that MYC activity amplifies the transcriptional induction of LAT1 under His limitation, although additional transcription factors, such as ATF4, are likely to contribute.

Together, these data demonstrate that His deprivation engages a conserved transcriptional program characterised by coordinated induction of amino acid transporters, even in cells with low basal MYC expression. ATF4 and MYC both contribute to this response, suggesting that His limitation elicits a stress-adaptive transcriptional network that promotes transporter expression and amino acid homeostasis independently of oncogenic MYC amplification.

### Histidine availability is a targetable metabolic vulnerability in glutamine-addicted MYC-driven tumours

Given the role of His in supporting LAT1-mediated amino acid exchange in Gln-dependent tumour cells, we next examined whether altering His availability influences tumour physiology *in vivo.* Because His is an essential amino acid, we first assessed the tolerability of dietary His restriction (HisFD) in healthy female FVB/N mice. Acute removal of dietary His resulted in rapid weight loss and reduced serum His levels (Fig. S7a,b), whereas His levels in peripheral tissues were only modestly affected (Fig. S7c), consistent with compensatory mechanisms that buffer EAA availability *in vivo*.

To enable longer-term studies while avoiding overt toxicity, we employed a diet containing 20%His (20%HRD). FVB/N mice were acclimated to control diet (CD) or 20%HRD prior to orthotopic injection of MYC-driven tumour cells (Fig. 7a). Tumour-bearing mice maintained on the 20%HRD exhibited reduced weight gain (Fig. 7b), accompanied by a significant reduction in serum and tumour His levels (Fig. 7c,d). Under these conditions, MYAZ16.1e tumours displayed significantly impaired growth, even after correcting for body-weight differences (Fig. 7e-g). Consistent with reduced amino acid availability, tumours from 20%HRD-fed mice exhibited decreased global protein synthesis, whereas liver tissue was insignificantly reduced (Fig. 7h and Fig. S7d). In line with our *in vitro* findings, His restriction increased MYC and Glyc and total ASCT2 levels, while LAT1 upregulation was observed in a subset of tumours (Fig. 7h), highlighting intra-tumoural heterogeneity in adaptive response.

**Fig. 7:**
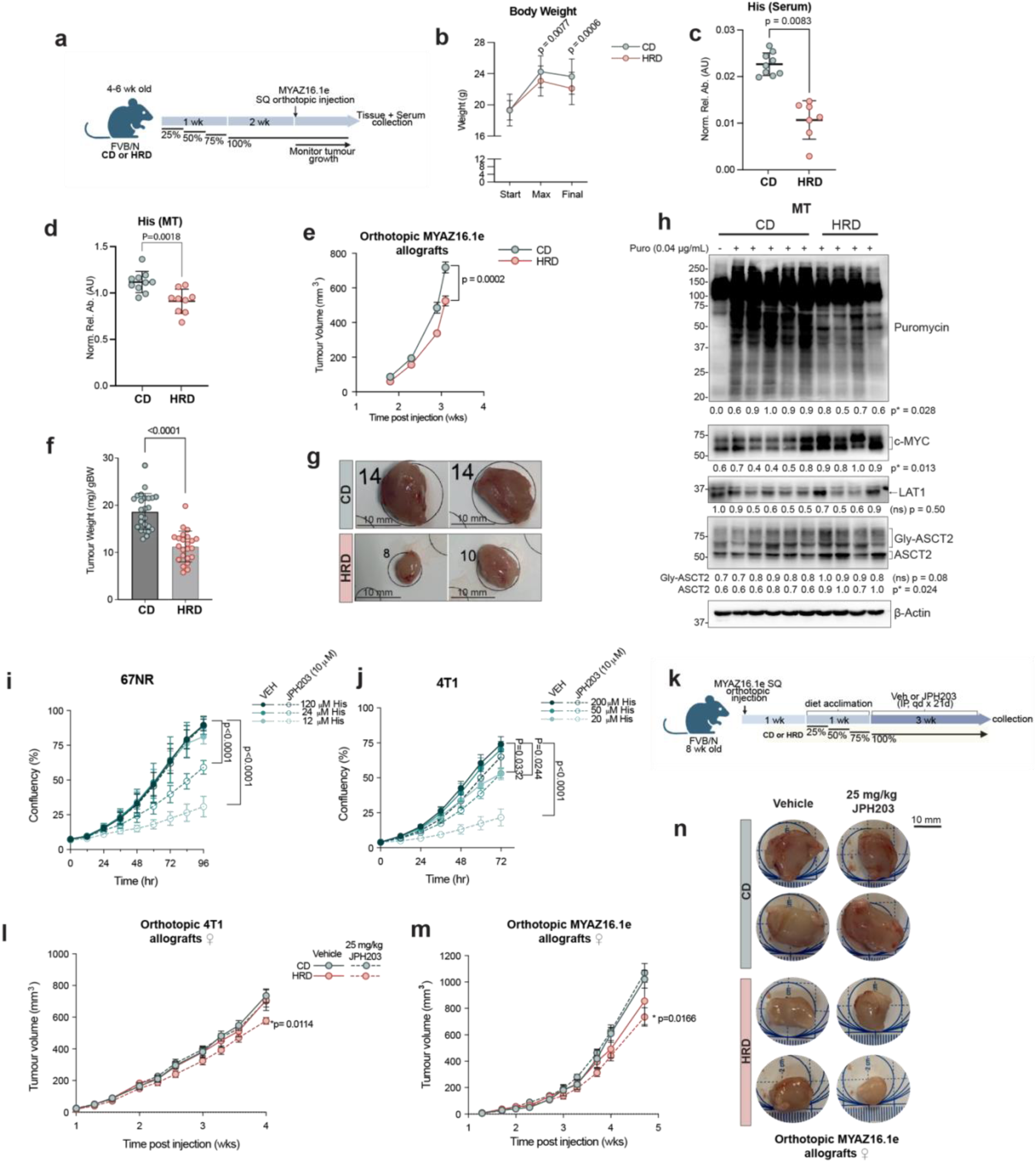
His availability is a metabolic vulnerability in glutamine-addicted MYC-driven tumours. (**a**) Experimental timeline: WT FVB/N mice on a control diet (CD) or 20%His-reduced diet (20%HRD) before subcutaneous (SQ) injection of MYAZ16.1e cells. (**b**) Body weight of CD-fed (n = 41) and 20%HRD-fed (n = 40) mice across two independent cohorts. Mean ± s.d.; *p*-values from two-way ANOVA (Tukey’s correction). (**c,d**) LC-MS analysis of (**c**) serum and (**d**) tumour His levels in CD (n = 9) and 20%HRD (n = 7) mice. Mean ± s.d.; *p*-values from unpaired two-tailed t-test. (**e**) Tumour volume in CD (n = 25) and 20%HRD (n = 25) mice. Mean ± s.d.; *p*-values from two-way ANOVA (Šídák’s correction). (**f**) Tumour weight normalized to body weight. Mean ± s.d.; *p*-values from unpaired two-tailed t-test. (**g**) Representative tumour images. (**h**) WB of MTs from subset of CD (n =5) and 20%HRD (n=4) mice injected with puromycin (Puro), plus an vehicle CD control. Protein levels were normalized to β-actin and expressed relative to the maximum. *p*-values from unpaired two-tailed t-test. (**i,j**) Proliferation of 67NR (**i**) and 4T1 (**j**) cells in indicated His concentrations ± JPH203 (n = 3). Mean ± s.d.; *p*-values from two-way ANOVA (Šídák’s correction). (**k**) Experimental timeline of tumour-bearing mice on a CD or 25%HRD ± JPH203. *IP: intraperitoneal, qd: quaque die, Veh: vehicle*. (**l,m**) Tumour volume of (**l**) 4T1: CD and Veh (n= 12), CD and JPH203 (n = 12), 25%HRD and Veh (n = 10), and 25%HRD and JPH203 (n = 11); and (**m**) MYAZ16.1e : CD and Veh (n= 8), CD and JPH203 (n = 6), 25%HRD and Veh (n = 7), and 25%HRD and JPH203 (n = 7) orthotopic allografts. Mean ± SEM; *p*-values from two-way ANOVA with Šídák’s correction. (**n**) Representative images of MYAZ16.1e tumours under each condition. **See also Figure S7, S8 and S9.**

We next asked whether His restriction enhances sensitivity to LAT1 inhibition. Under His-replete conditions, c-*Myc*-low 67NR cells were largely insensitive to the LAT1 inhibitor, JPH203, and His limitation alone had no effect on proliferation (Fig. 7i). In contrast, combining His limitation with JPH203 treatment resulted in a pronounced suppression of cell growth, revealing a synthetic lethal interaction between reduced His availability and LAT1 inhibition in this otherwise resistant cell line (Fig. 7i). Consistent with this, His limitation decreased the IC_50_ of JPH203 from 24.4 µM to 8.9 µM (Fig. S8a). By comparison, c-*Myc*-high 4T1 cells exhibited greater baseline sensitivity to both His limitation and LAT1 inhibition, consistent with increased reliance on amino acid transport (Fig. 7j; Fig. S8b). Nevertheless, combined His limitation and JPH203 treatment further exacerbated growth suppression in these cells (Fig. 7j), indicating that His restriction enhances sensitivity to LAT1 inhibition in tumour cells with differing MYC status.

To assess whether this interaction could be recapitulated *in vivo*, we next tested dietary His restriction in combination with LAT1 inhibition. To mitigate the reduced weight gain observed at 20%HRD and enable combination treatment, we employed a milder 25% histidine-reduced diet (25%HRD). In healthy mice, 25%HRD lowered serum His levels over an 8-week period (Fig. S9a) and modestly reduced weight gain without affecting food intake (Fig. S9b,c). Importantly, 25%HRD did not induce muscle atrophy, as indicated by preserved muscle mass and unchanged serum and muscle 3-methylhistidine levels (Fig. S9d), and instead preferentially reduced fat mass (Fig. S9f).

Using this regimen, orthotopic 4T1 and MYAZ16.1e tumours were first established and mice were subsequently transitioned to CD or 25%HRD prior to treatment with JPH203 (Fig. 7k). Consistent with previous reports, JPH203 alone had no effect on tumour burden in CD-fed mice in either model. While 25%HRD alone produced a modest reduction in tumour growth in MYAZ16.1e allografts, the combination of His restriction and LAT1 inhibition resulted in a statistically significant reduction in tumour burden in both models (Fig. 7l-n). These data indicate that limiting His availability enhances the tumour growth-suppressive effects of LAT1 inhibition *in vivo*.

In summary, our findings define a mechanism by which Gln-dependent tumours, including MTs induced by MYC, exploit His to sustain amino acid homeostasis and growth. MYC-hyperactivation upregulates amino acid transporters, including ASCT2 and LAT1, to support anabolic metabolism and protein synthesis. In this context, Gln-addicted cells preferentially utilise glitamine for anaplerosis, which may limit its availability for LAT1-mediated EAA exchange. Given that the tumour microenvironment is Gln-poor yet relatively rich in His, intracellular His preferentially facilitates LAT1-dependent uptake of EAAs, sustaining mTORC1/4E-BP1 signaling and translation. Under His-limited conditions, MYC- and ATF4-dependent induction of ASCT2 and LAT1 partially compensate for amino acid loss but concurrently increases tumour sensitivity to LAT1 inhibition. Together, these findings identify His availability as a physiological constraint on amino acid exchange that can be exploited to sensitize Gln-dependent tumours to LAT1 inhibition *in vivo*.

## Discussion

Our results identify histidine as a major LAT1 exchange substrate in glutamine-dependent MTs, where it supports mTORC1/4E-BP1 signaling, amino acid homeostasis, and tumour growth. This refines the prevailing model that glutamine is the principal LAT1 exchange factor driving mTORC1 activation^1^. The preferential use of histidine, which is present at comparable levels to Gln in the TME, may reflect its minimal catabolism and high affinity for LAT1. These observations suggest that LAT1 substrate utilisation is context-dependent and adapts to tumour-specific metabolic demands, with His representing a more cost-effective choice in glutaminolytic cancer cells.

Mechanistically, our data indicate that LAT1 predominantly mediates histidine efflux while contributing partially to its uptake. Recent work has identified SNAT3 (encoded by SLC38A3) as a His transporter in cancer cells^24^, and we observe that SLC38A3 expression is co-upregulated with SLC7A5 during MYC-driven tumourigenesis (Fig. 1a,b). Consistently, MYC-driven MTs exhibit increased His uptake and accumulation compared to the NMG. Elevated histidine availability selectively potentiates mTORC1/4E-BP1 signaling over the mTORC1/S6K1/RPS6 axis in a LAT1-dependent manner. In line with this, MYC/eIF4E-dependent neural clones displayed heightened sensitivity to His deprivation relative to S6K-dependent clones in a previous study^25^.

Unexpectedly, histidine limitation increased both c-MYC transcription and translation, in contrast to reports that LAT1 ablation suppresses MYC translation and protein abundance^12^. The mechanisms linking His availability to MYC transcription remain unclear and warrant further investigation.

Previous work has shown that *SLC7A5* knockdown alone does not impair proliferation under amino acid-replete conditions, whereas limiting high-affinity LAT1 substrates exacerbates growth defects^24^. In agreement, we find that MYC-low cells become sensitive to LAT1 inhibition only under His-restricted conditions. Conversely, MYC-high MT cells are intrinsically vulnerable to both His deprivation and LAT1 inhibition. Histidine limitation robustly induces *Slc7a5*/LAT1 expression via MYC and ATF4 co-regulation, reinforcing his availability as a critical vulnerability in MYC-driven tumour cells. Whether His restriction elicits a distinct transcriptional programme or amplifies canonical amino acid stress responses remains to be determined.

Finally, we demonstrate that systemic modulation of His availability impacts tumour progression. A His-reduced diet suppresses tumour growth while selectively decreasing fat mass and preserving lean mass, consistent with other EAA restriction strategies^26,27^. Although the LAT1 inhibitor JPH203 shows limited efficacy as a monotherapy in MYC-driven MTs, HRD significantly, albeit modestly, enhances tumour sensitivity to LAT1 inhibition, supporting a combinatorial therapeutic approach.

Collectively, our study establishes histidine availability as a key determinant of LAT1 function and tumour metabolism in glutamine-dependent cancer cells. These findings highlight the potential of targeting nutrient availability to modulate transporter activity and exploit metabolic dependencies. More broadly, they provide a conceptual framework for leveraging dietary interventions to regulate amino acid transport and improve therapeutic outcomes in highly glutaminolytic tumours.

## Methods

### Cell Culture

#### Generation of MYC-driven tumour cell lines

MYC-driven tumour cell lines (MYAZ16.1e, MYAZ6.6g, and MYAZ12.3i) were established from MMTV-c-*Myc* MTs. Tumours were minced on ice using sterile razor blades and enzymatically dissociated in Digestion Buffer (MMEC media supplemented with 1% BSA, 1 mg/mL Collagenase Type L) at 37°C for 30 min at 200 rpm. The resulting suspension was passed through a 70 µm filter, centrifuged (530 *xg*, 5 min), and resuspended in 5 mL RBC lysis solution (170 mM Tris, 150 mM ammonium chloride, pH 7.4) before a final centrifugation step. Cells were cultured in MMEC media (DMEM supplemented with 10% heat inactivated FBS (iFBS), 10 mM HEPES, 2 mM Gln, 5 µg/mL insulin, 1 µg/mL hydrocortisone, 10 ng/mL EGF, and 100 µg/mL Penicillin/Streptomycin) to establish tumour cell lines.

#### Culture Conditions and Authentication

Additional cell lines used included 67NR, 4T1, BT-549, Hs578T, T47D, AU565, and HEK293T. Human cell lines were authenticated by STR (short tandem repeat) profiling. All cell lines (mouse and human) were routinely tested for mycoplasma contamination and confirmed negative. Tumour cell lines were maintained in MMEC media, while all other cell lines were cultured in DMEM (Gibco) supplemented with 10% iFBS, 2 mM Gln (Gibco), and 1% Penicillin-Streptomycin. All cell lines were maintained under standard conditions (37°C and 5% CO_2_ , atmospheric O_2_).

#### Experimental DMEM Preparation

Custom DMEM formulations were prepared by the Francis Crick Institute Media Preparation Service based on DMEM (Cat No. 11960044, Gibco) and included: (i) amino acid-free DMEM, (ii) His, Ile, Met, Leu, and Gln-free DMEM, (iii) His and Gln-free DMEM, and (iv) Gln and glucose-free DMEM. L-His, L-Ile, L-Met, L-Leu, L-Gln, and D-glucose were supplemented at the indicated concentrations. Where specified, homocysteine, vitamin B_12_, D-His, and doxycycline were added.

#### Preparation of Dialysed Serum

For experiments requiring amino acid-free or reduced-amino acid conditions, dialysed iFBS was prepared by dialyzing FBS against calcium- and magnesium-free PBS (137 mM NaCl, 2.7 mM KCl, 10 mM Na_2_HPO_4_, 1.8 mM KH_2_PO_4_, pH 7.0) using dialysis tubing (3,500 Da MWCO, Cat No. 68035, ThermoFisher Scientific). PBS was changed daily for 5 days.

#### Generating stable ATF4 knockdown cells

Stable ATF4 knockdown in 67NR cells was achieved via lentiviral transduction. Lentiviral particles were produced by transiently transfecting HEK293T cells in 60-mm culture dishes with 2 µg shRNA-expressing vectors (pLKO-puro sh*Atf4* or sh*Luc*, kindly provided by Dr. Andrés Méndez Lucas, Universitat de Barcelona), 750 ng psPAX2, and 250 ng pMD2.G.

Transient transfection was performed using polyethylenimine (PEI, 10 µg/mL) at a 3:1 PEI:DNA ratio. After 16 h, the medium was replaced, and viral supernatants were collected at 48 h post-transfection, centrifuged (1,000 x*g*, 5 min), and filtered through a 0.45 µm PES filter.

For infection, 67NR cells were incubated with viral supernatant supplemented with 8 µg/mL polybrene for 48 h. Selection was performed using 1 µg/mL puromycin.

Stable multi-clonal cultures were propagated for experiments, and knockdown efficiency was confirmed via RT-qPCR.

### CRISPR Cas9 genome editing

#### sgRNA design and cloning

CRISPR/Cas9 target sites for *SLC7A5* and c-*Myc* were identified using the CHOPCHOP web tool (https://chopchop.cbu.uib.no/). 24-bp oligonucleotides were synthesized with a 4-bp overhang, where the forward oligo contained a TCCC overhand, and the reverse complement oligo contained an AAAC overhang for cloning in the BsmB1 site of FGH1tUTG (Addgene #70183, a gift from Marco Herold). Successful clones were sequence-verified.

sg*SLC7A5*: (F) 5’-TCCCTGTCTCCACAGTGCTGCTCA, (R)5’-AAACTGAGCAGCACTGTGGAGACA.

sg*Myc*:(F) 5’-TCCCTCGGGCTCATCTCCATCCCG, (R) 5’-AAACCGGGATGGAGATGAGCCCGA

#### Lentiviral gene transfer, clonal selection, and validation

Lentiviral particles were produced in HEK293T cells via transient transfection, as described above. Stable *SLC7A5* or c-*Myc* knockout in BT-549 and MYAZ16.1e cells, respectively, was achieved using a two-vector CRISPR-Cas9 system: Cas9 expression via FuCas9Cherry (Addgene #70182) and sgRNA expression via Fgh1tUTG (Addgene #70183), which provides Dox-inducible sgRNA expression and includes a GFP reporter. Following lentiviral transduction, double-positive (mCherry^+^ GFP^+^) cells were selected. For BT-549 sg*SLC7A5* cells, a multi-clonal culture was generated by sorting double-positive cells. For MYAZ16.1e sg*Myc* cells were sorted as single cells into 96-well plates using the MoFlo XDP cell sorter (Beckman Coulter) to generate clonal populations. Successful knockouts were confirmed by immunoblotting.

#### Growth Assays

For proliferation assays, cells were washed with PBS before being seeded in 96-well plates under indicated experimental conditions. Cells were imaged at regular intervals using the Incucyte^®^ live-cell imaging system, and confluency was quantified using the Incucyte S3 Software (Sartorius v.2021C).

#### SUnSET Assay

Nascent global protein synthesis was measured using the SUnSET (Surface Sensing of Translation) assay^28^.

##### in vitro

Cells were treated with 10 µg/mL puromycin for 10 min before collection for lysate preparation. Following treatment, cells were washed with ice-cold PBS on ice and lysed as described below for western blot (WB) analysis.

##### in vivo

Mice were administered an intraperitoneal (IP) injection of 0.02 mg/g body weight (BW) puromycin (5 mg/mL in 0.9% saline). After 30 min, tissues were rapidly dissected and flash-frozen in liquid nitrogen for subsequent WB analysis.

#### Preloading Assays

MYAZ16.1e and BT-549 cells were cultured in amino-acid free medium (-AA) supplemented with 2% dFBS, with or without the indicated concentrations of His, Gln, or Asn (preload) for 48 h. A control plate (Ctl) was collected before amino acid starvation to assess baseline amino acid levels and mTORC1 activation. After 48 hours, cells were either collected immediately (-AA or -AA+H) or washed with PBS at room temperature and replenished with His-, Gln-, or Asn-free DMEM for the indicated duration (-AA/DMEM or -AA+X/DMEM). Cells and media were then collected and flash-frozen in liquid nitrogen for subsequent metabolite extraction and LC-MS analysis or protein extraction for WB analysis.

#### Stable isotope labelling *in vitro*

Unlabelled amino acids were replaced with their isotope-labelled counterparts, [U^13^C]His and [U-^13^C]Gln (Cambridge Isotope Laboratories, Inc.), in DMEM at the indicated concentrations and durations.

#### Quantitative real-time RT-PCR (qPCR)

RNA extraction and qPCR were performed as previously described^29^. Briefly, cells were collected in TRIzol™ (ThermoFisher Scientific), while tumour tissues (∼ 3-5 mg) were homogenized in TRIzol™ for RNA extraction following the manufacturer’s protocol. RNA quality and concentration was assessed using a NanoDrop™ 2000 spectrophotometer (ThermoFisher Scientific). Reverse transcription was performed using Superscript™ III Reverse Transcriptase (ThermoFisher Scientific). Quantitative PCR was performed using Power SYBR™ Green PCR Master Mix (ThermoFisher Scientific) on a ViiA 7 Real-Time PCR system (Applied Biosystems) according to the manufacturer’s protocol. Relative gene expression was determined using the 2^-Δ^ ^Δ^*^Ct^* method, normalized to β-actin.

The primers used for qPCR were as follows: *c-Myc* (forward: TTGGAAACCCCGCAGACA, reverse: CGGAGTCGTAGTCGAGGTCA), *Slc7a5* (forward: CTGGTCTTCGCCACCTACTT, reverse: GCCTTTACGCTGTAGCAGTTC) was from Poncet et al. (2014)^30^, the following primers were from Bian et al. (2020)^31^: *Slc7a6* (forward: TCTACCTTCGCTGGAAAGAGCC, reverse: GCCACCAGAAACAAGGAGCAGA), *Slc7a7* (forward: AAGGTGTTGGCGCTGATTGCAG, reverse: AGAGTGCCAGAGCAATGTCACC), *Slc7a8*(forward: GCATACGTCACTGCAATGTCCC, reverse: GGAGCCATTGACTCCACCAAAC), *Slc38a1* (forward: TACCAGAGCACAGGCGACATTC, reverse: ATGGCGGCACAGGTGGAACTTT), *Slc38a2* (forward: GCGTTGGCATTCAATAGCACCG, reverse: TCGTAGATGGGAAGAACAGCGG), *Slc43a1* (forward: TTCCTGTGGAGCCTTGTCACCA, reverse: CTCCACCTTCTGTCTCTGCTCA), *Slc43a2*(forward: CAGCATCCTTGAGTTCCTGGTC, reverse: TGATGTAGCCGATGACAGGAGC), and *Slc1a5* (forward: GCCTTCCGCTCTTTTGCTAC, reverse: GACGATAGCGAAGACCACCA), *ActB* (forward: GGCTGTATTCCCCTCCATCG, reverse: CCAGTTGGTAACAATGCCATGT) were from Kreuzaler et al. (2023)^29^ . *Slc7a1* (forward: GCAAAAACCTGCTCGGTCTG, reverse: TCATAGGTGTTGAGGCAGCG), *Slc36a1* (forward: GGTCGGGAGAGGTAGAGGTT, reverse: AAAGAGCAGACACTGACGGG), *Atf4* (forward: TCCCTTTCCTCTTCCCCTCC, reverse: GGATTTCGTGAAGAGCGCCA), and *Slc38a3* (forward: CTACGAGCAGTTGGGCTACC, reverse: TGCCATCCATGTACCACACC), *Slc38a5* (forward: TGAATGGAACCCTCTCTGCG, reverse: CCCCATGATAGCGTTGCTGA), *Slc38a8* (forward: ATCCTCTTGAAGTCCGCTCTG, reverse: GCATGCCTCCTGCCTTGTAG) were designed using NCBI Primer-BLAST (https://www.ncbi.nlm.nih.gov/tools/primer-blast/).

#### RNA-seq Analysis

RNA sequencing data were analyzed by the Bioinformatics & Biostatistics Science Technology Platform at the Francis Crick Institute. RNA-sequencing reads were processed using the nf-core RNA-seq pipeline (version 3.3) with the STAR/RSEM option. Reads were aligned to the mouse genome assembly GRCm38 using Ensembl release 95 transcript annotations. Gene-level abundance estimates were obtained from RSEM and imported into R using the DeSeqDataSetFromMatrix function in DESeq2. Normalization factors were computed per gene, correcting for library size and feature length. Variance stabilizing transformation (VST) was applied to abundance estimates to standardize gene expression across samples. Principal component analysis (PCA) was performed using all detected genes. Hierarchical clustering based on Euclidean distances was used to generate heatmaps of sample relationships.

RNA-seq data from MMTV-c-*Myc* derived MT cell lines was analyzed in R (version 2024.04.2). Gene expression counts were filtered for genes of interest, annotated via Ensembl (biomaRt version 2.58.2), and Z-score normalized across samples. Hierarchical clustering and heatmap generation was performed using pheatmap (version 1.0.12)

Publicly available gene expression data (<u>GSE75957</u>)^13^ were analyzed for His metabolism-related genes in wildtype FVB/n adult female tissues. Data were filtered in R (version 2024.04.2) to include relevant samples and genes, Z-score normalized RPSM values were used, and hierarchical clustering was applied. A heatmap was generated using the pheatmap to visualise gene expression patterns.

#### Western Blotting

Cells were lysed in Triton-X lysis buffer (1% Triton X-100, 150 mM NaCl, 50 mM Tris, pH 8.0) supplemented with protease and phosphatase inhibitors. Tissues were homogenized in the same buffer using a Precellys Cryolys Evolution tissue homogenizer (Bertin Technologies). Lysates were incubated on ice, centrifuged (21,000 x*g*, 15 min, 4°C), and protein concentrations were determined using the Pierce™ BCA Protein Assay Kit (ThermoFisher Scientific). Samples were prepared in Laemmli buffer supplemented with 10% 2-mercaptoethanol, and heated at 37°C. For histone extraction, nuclei were isolated by resuspending cell pellets in Triton X-100 lysis buffer, followed by sequential centrifugation and washing steps. Histones were extracted in 0.2 N HCl, sonicated and incubated overnight at 4°C before neutralization with NaOH. Protein concentration was measured using the Pierce™ BCA Protein Assay Kit, and samples were prepared in Laemmli buffer as above.

For immunoblotting, 10-20 µg of protein was resolved by SDS-PAGE (10-12% gels) and transferred onto nitrocellulose membranes via wet transfer at 400 mA for 1-2 h at 4°C. Membranes were blocked in either 5% BSA (for phospho-proteins) or 5% non-fat dry milk in TBST, incubated with primary antibodies in 5% BSA in TBST overnight at 4°C, washed, and probed with HRP-conjugated secondary antibodies. Detection was performed using ECL reagent (Cytiva Life Sciences) and an Amersham ImageQuant™ 800 imager. For reprobing, blots were stripped using a glycine-SDS buffer (0.2 M glycine, 1% SDS, 0.1% Tween-20, pH 2.2), re-blocked, and incubated with additional antibodies. Protein quantification was performed using FIJI (https://github.com/fiji), normalizing protein levels to loading controls and, where applicable, to total un-phosphorylated protein levels.

#### Immunofluorescence

Cells grown on glass coverslips in indicated experimental conditions were washed three times with ice-cold PBS and fixed in 4% paraformaldehyde (PFA) for 12 min at room temperature. Fixed cells were washed five times with PBS before blocking and permeabilization with 3% BSA and 0.2% Triton X-100 in PBS for 12 min at room temperature. After three PBS washes, cells were incubated overnight at 4°C with primary antibodies diluted in 3% BSA in PBS. The primary antibodies used were ASCT2 (1:200; Santa Cruz Biotechnology).

Following incubation, cells were washed five times with PBS and incubated with fluorescently conjugated secondary antibodies in 3% BSA in PBS for 1 h at room temperature in the dark. The secondary antibodies used were Alexa Fluor 488 anti-rabbit (Invitrogen, A-11008, 1:400). After five additional PBS washes, nuclei were stained with DAPI (1:10,000) for 5 min, followed by another five PBS washes. Coverslips were mounted using ProLong™ Gold Antifade Mountant (ThermoFisher Scientific).Imaging was performed using an Axio-Imager M2 upright microscope (ZEISS), and images were processed with FIJI.

#### Antibodies

The following primary anitbodies were used in this study: anti-c-MYC Y69 (Abcam, ab32072; 1:1000), anti-ATF4 (Abcam, ab23760; 1:1000), anti-p-eIF2𝘢 (Ser51) (Cell Signaling Technology, 3398; 1:1000), anti-eif2𝘢 (Cell Signaling Technology, 5324; 1:1000), anti-ASCT2 (Santa Cruz Biotechnology, sc-99002; 1:1000), anti-P-4E-BP1 (Ser65) (Cell Signaling Technology, 9451; 1:1000), anti-4E-BP1 (Cell Signaling Technology, 9644; 1:2000), anti-P-RPS6 (Ser 240/244) (Cell Signaling Technology, 4858; 1:1000), anti-RPS6 (Cell Signaling technology, 2317; 1:2000), anti-Gls1 (Abcam, ab93434; 1:1000), anti-Gls2 (ProSci Inc., 6217; 1:1000), anti-Glutamine Synthetase (BD Biosciences, 610518; 1:1000), anti-H3K4me (Abcam, ab8995; 1:1000), anti-H3K4me3 (Merck, 05-745R; 1:1000), anti-HER2/ErbB2 (Abcam, ab16901; 1:1000), anti-LAT1 (human, Cell Signaling Technology, 5347; 1:1000), anti-LAT1 (2B Scientific, KAL-KE026; 1:300), anti-LC3B (Cell Signaling Technology, 43566; 1:1000), anti-mTOR (Cell Signaling Technology, 2972; 1:1000), anti-Puromycin (Merck, MABE343; 1:1000), anti-omoMYC (Cancertools, 153657; 1:1000) anti-Vinculin (Abcam, ab129002; 1:1000) β-actin (HRP-coupled, Merck, A3854; 1:25,000)

#### Mice

##### Husbandry

All mouse procedures and husbandry complied with United Kingdom Home Office regulations under the Animals (Scientific Procedures) Act 1986 and were approved by the Animal Welfare and Ethical Review Panel under project license PP3464389 at the Francis Crick Institute.

Mice were bred and housed in groups of up to five per cage under specific-pathogen-free conditions at the Francis Crick Institute animal facility. All mice had *ad libitum* access to chow diet (Envigo) and water.

##### Mouse Models

This study utilized wildtype FVB/NJ and Balb/c females from in-house colonies at the Francis Crick Institute. Additionally, transgenic MMTV-c-*Myc* (Tg(MMTV-MYC)141-3Led, MGI ID: 2447500) mouse model backcrossed onto the FVB/NJ background for at least 10 generations was included, originally obtained from The Jackson Laboratory. Transgenic mice were genotyped using PCR-based assays.

##### Orthotopic tumour generation

Orthotopic mammary tumour allografts of MYAZ16.1e and 4T1 cells were generated by injecting a Matrigel™-cell suspension into the fourth inguinal mammary fat pad of 8- to 10-week-old virgin FVB/NJ or Balb/c mice, respectively. Tumours were monitored 2-3 times weekly for growth and signs of distress. Humane endpoints were defined as 20% total body weight loss or 15% weight loss within 72 h. Tumour dimensions were measured using calipers, and volume was estimated using the standard ellipsoid formula.

##### Diet modifications and compound administration

Mice were placed on synthetic amino acid-defined diets (TestDiet^®^ Baker AA, IPS Product Supplies Ltd.) to modulate His availability. For His restriction, diets included a no-added His diet (5WTU), a 25% His diet (5GEG), a 20% His diet (5GFQ), and a His control diet (5WA1). Other amino acids were proportionally adjusted to maintain total amino acid intake.

For experiments in which the diet was provided before orthotopic injection of MYAZ16.1e cells, mice were transitioned onto His-modified diets (5GFQ and 5WA1) at 5- to 6-weeks of age over six days and remained on their assigned diet for three weeks before tumour cell injection. Mice continued their respective diets until the experimental endpoint. For experiments in which the diet was introduced after tumours became palpable, mice were transitioned onto His-modified diets (5GEG and 5WA1) over six days and maintained on their assigned diet in combination with either vehicle or 25 mg/kg JPH203 (dissolved in 40% (*w/v*) 2-hydroxypropyl-β-cyclodextrin (HP-β-CD) in 0.9% (*w/v*) saline; S866705, Selleck Chem) via daily intraperitoneal (IP) injection for up to 21 days.

##### Metabolic Cages for Food Intake

Wildtype 8-week-old FVB/N mice were transitioned onto His-modified diets (5GEG and 5WA1) over six days, after which they remained on their assigned diets. Mice were then individually housed in metabolic cages (Animalab) and acclimated for four days. Following acclimation, a pre-weighed amount of the assigned diet was provided daily, and the remaining food was weighed over a 7-day period to monitor food intake.

##### Deuterium Oxide Administration to Estimate Body Composition

Mice were administered deuterium oxide (^2^H_2_O, Cambridge Isotope Laboratories) to achieve a target enrichment of 3% of total body water (calculated as 70% of body weight). A stock solution of ^2^H_2_O in 0.9% (*w/v*) saline was prepared, and mice were injected intraperitoneally with a bolus. To maintain enrichment, drinking water was supplemented with ^2^H_2_O at a final concentration of 3% (*v/v*). Blood was collected 24 h post-injection. Serum was isolated and duplicate samples were denatured using acetone. Samples were centrifuged (21,000 x*g*, 15 min, 4°C), and the resulting supernatant was transferred to 3 mm NMR tubes for analysis. ^2^H NMR measurements were acquired through 1D pulse-and-acquire. ^2^H NMR spectra were recorded at 25°C on a 700 MHz Bruker Avance III NMR spectrometer equipped with a 5 mm QCl cryoprobe with no field-frequency lock. A 30° 2H pulse was employed, the relaxation delay was 8 s and the acquisition time was 1.6 s (sweepwidth 20 ppm; 6880 complex points). The number of transients was 64, with 4 dummy scans. The free induction decays were apodized with 4 Hz line broadening and zero-filled to 32K points, prior to Fourier transformation, baseline correction and signal integration, all conducted in TopSpin (version 3.6.4; Bruker). The integral of the ^2^H_2_O and acetone peaks was calculated and fitted to a calibration curve to determine ^2^H_2_O enrichment in serum. Total body water (TBW) was calculated using the dilution principle, dividing the amount of administered ^2^H_2_O by its concentration in the serum. Fat free mass (FFM) was estimated as previously described^32^, based on the assumption that lean tissue contains ∼73% water. Fat mass was determined by subtracting the FFM from the total body weight.

#### Stable isotope labelling *in vivo*

Stable isotope labelling was performed using either a bolus injection or a continuous infusion to assess amino acid uptake or metabolic incorporation. Short-term boluses of [U^13^C]His were administered intravenously to evaluate tissue uptake, while a longer 3 h infusion of [U^13^C]His was used to track metabolic incorporation in wildtype FVB/N adults and tumour bearing transgenic MMTV-c-*Myc* mice. For bolus injections, [U^13^C]His (Cambridge Isotope Laboratories) was diluted to 25 mg/mL in 0.9% saline and administered at 0.107 mg/gBW for 5 min. For His infusions, [U^13^C]His was delivered via a tail vein catheter using an Aladdin AL-1000 pump (World Precision Instruments) in mice under isoflurane anaesthesia. A priming bolus of 0.12 mg/gBW was first administered, followed by a 3.45 µg/gBW/min continuous infusion over 3 h. At the end of each bolus or infusion, blood was collected via cardiac puncture, allowed to clot on ice, centrifuged (10,000 rpm, 10 min, 4°C), and serum was flash-frozen in liquid nitrogen. Tissue was flash-frozen in liquid nitrogen.

#### Immunohistochemistry

Tissues were placed in 10% neutral buffered formalin, incubated for 48 h at 4°C with rotation, and then transferred to 70% ethanol for storage. Tissues were processed through graded alcohols then Xylene and finally embedded in paraffin wax. Sections of 3 µm were cut for downstream staining. Hematoxylin and Eosin (H&E) staining was performed using a Tissue-Tek Prisma Plus Automated Slide Stainer & Coverslipper (Sakura Finetek). Following deparaffinization and rehydration, the sections were stained with Hematoxylin for 5 min, rinsed with water, before incubation in 1% acid alcohol for 20 s. After another water wash and incubation in Scott’s tap water for 1 min, the sections were counterstained with Eosin for 5 min. Finally, they were dehydrated using a graded series of ethanol and cleared with xylene.

For IHC, sections were baked for 1 h at 60°C before IHC staining was performed using the Leica Bond-Rx autostainer platform. Sections were stained with anti-LAT1 antibody [1:75, CosmoBio, KAL-KE026] for 15 min at RT. Epitope antigen retrieval solution 2 (AR9640, Leica) was used for 20 min at 95°C prior to the antibody application. BOND Polymer Detection kit (DS9800, Leica) was used, including hydrogen peroxidase, anti-rabbit polymer, DAB, and haematoxylin counterstain. Slides were counterstained with haematoxylin then coverslipped using Tissue -Tek Glass™ g2 Automated Glass Coverslipper. Images were acquired using Axioscan 7 brightfield microscope slide scanner (ZEISS) at 20x magnification. Images were processed using QuPath (version 0.5.1). LAT1 IHC membrane staining intensity was quantified using a previously published method^33^. Briefly, a custom Groovy script was used for cell segmentation, utilizing a watershed-based cell and membrane detection algorithm. Image deconvolution was applied to isolate the membrane compartment. Staining intensity was categorized as high (> 1.0), medium (>0.6 to ≤ 1.0), or low (>0.3 to ≤ 0.6) based on mean DAB signal intensity. The quantification provided the total number of cells and the number classified into each staining category.

#### Metabolomics

##### Tissue Extraction

Snap-frozen tissues were ground in liquid nitrogen using a mortar and pestle, then lyophilized overnight in a FreeZone 4.5 Freeze Dry System (Labconco). Notably, the NMG tissue was not lyophilized due to its high fat content.

Polar metabolites were extracted using biphasic extraction. For each sample, 3-5 mg of lyophilized tissue was extracted with 1.2 mL chloroform and 0.6 mL of methanol (2:1) containing the internal standard, 5 µM [final] [U^13^C^15^N]Valine. Samples were vortexed at maximum speed for 30 s each, sonicated for three rounds of 8 min each in a water bath sonicator at 4°C over 1 h. Following this, samples were centrifuged (21,000 x*g*, 20 min, 4°C), and the supernatant (SN1) was collected and vacuum-dried using a rotational-vacuum-concentrator RVC 2-33 CD (Christ). The remaining pellet was re-extracted with 0.5 mL methanol and 0.25 mL water (2:1). After vortexing and sonication as above, the samples were centrifuged (21,000 x*g*, 20 min, 4°C). The resulting supernatant (SN2) was combined with SN1 and vacuum dried. The final pellet was retained for protein quantification.

Once the samples were thoroughly dried, biphasic partitioning was performed to separate polar and apolar phases. A 1:3:3 ratio of chloroform:methanol;water was added to each sample, followed by vortexing for 30 s. Samples were then centrifuged (21,000 x*g*, 5 min, 4°C), resulting in two distinct phases. The upper polar phase was analysed by LC-MS; if not immediately submitted for analysis, the samples were vacuum-dried and stored at -80°C.

##### Cell Extraction

Cells were seeded in a 6-well plate for all metabolomics experiments under the indicated experimental conditions. At collection, cells were rinsed 1x with room temperature PBS before submerging the plate in liquid nitrogen and maintaining it on dry ice until metabolite extraction.

Cells were scraped with 0.75 mL ice-cold methanol, 0.25 mL water with 5 µM [final] [U^13^C^15^N]Valine as the internal standard, and 0.25 mL chloroform (3:1:1). Samples were vortexed and sonicated as above, and incubated overnight at 4°C.

After incubation, samples were centrifuged (21,000 x*g*, 10 min, 4°C), and the supernatant was collected, the pellet was retained for protein quantification. 0.5 mL of water was added, adjusting the methanol:water:chloroform ratio to 3:3:1, to enable biphasic partitioning. The samples were vortexed and centrifuged (21,000 x*g*, 5 min, 4°C). The upper polar phase was vacuum-dried using a rotational-vacuum-concentrator RVC 2-33 CD (Christ) at 30°C. The vacuum-dried polar phase was either stored at -80°C or resuspended in 100-200 µL of 1:1 mixture of methanol:water, and 100 µL was analyzed by LC-MS.

##### Media and Serum Extraction

Media from *in vitro* experiments and serum collected from blood were extracted following the same protocol. For each sample, 10-15 µL was first precipitated with a 3:1 ratio of ice-cold methanol. Samples were vortexed and incubated on ice for 5 min to allow precipitation, followed by centrifugation (21,000 x*g*, 10 min, 4°C).

A 1:1 mixture of methanol and water was prepared with 5 µM [final] [U^13^C^15^N]Valine. The supernatant was collected for biphasic partitioning using a 3:3:1 ratio of methanol:water:chloroform. The samples were vortexed and centrifuged (21,000 x*g*, 5 min, 4°C), yielding two phases. The upper polar phase was isolated and submitted for LC-MS analysis.

##### Protein quantification for normalization

The pellets from methanol-chloroform extractions were used for protein quantification to normalize LC-MS data where dry weight was not available. For experimental subsets with tissues where the NMG was not included in the analysis, the dry weight was used for normalization.

Pellets were resuspended in 100 µL of freshly prepared Urea lysis buffer (6M Urea, 15 mM DTT, 2% CHAPS). Samples were vortexed for 30 s, sonicated for 8 min at 4°C, and incubated at room temperature on an orbital shaker at 100 rpm for 30 min. After incubation, samples were centrifuged (10,000 x*g*, 5 min, room temperature).

BSA standards were prepared in the urea lysis buffer, and the Pierce™ 660 nm protein assay kit (ThermoFisher Scientific), was used for protein quantification according to manufacturer’s instructions.

##### Absolute quantification of Gln and His by LC-MS

To quantify absolute Gln and His concentrations in serum and TIF, a standard curve of [U^13^C]His and [U^13^C]Gln was generated, ranging from 750 nM to 37.5 µM final concentration. Stable isotope-labelled internal standards ([U^13^C]His and [U^13^C]Gln) were spiked into all samples at a known concentration prior to extraction to correct for matrix effects.

##### LC-MS Instrumentation

Polar extracts were analysed by LC-MS using a Q-Exactive Plus (Orbitrap) Mass Spectrometer coupled to a Vanquish UHPLC system (ThermoFisher Scientific), as previously described^34^. Chromatographic separation of polar metabolites was achieved using a SeQuant ZIC-pHILIC column (150 mm x 4.6 mm, 5 µm particle size; Merck) at a flow rate of 300 µL/min. A 15-minute gradient elution was applied, transitioning from 80% solvent A (20 mM ammonium carbonate in water, Optima LC/MS grade) to 20% solvent B (acetonitrile, Optima LC/MS grade), followed by a 5 min wash at 95:5 (solvent A : solvent B) and a 5 min re-equilibration. The injection volume was 5 or 10 µL as appropriate, and the autosampler was maintained at 4°C. The mass spectrometer operated in positive/negative polarity switching mode, scanning across a mass range of 70-1050 m/z with a resolution of 70,000. Ionization was performed using a heated electrospray ionization (HESI-II) probe with a spray voltage of 3.5 kV (positive mode) and 3.2 kV (negative mode). Additional MS parameters included a probe temperature of 320°C, sheath gas and auxiliary gas settings of 30 and 5 arbitrary units, and an AGC target of 3 x 10^6^. Prior to analysis, mass calibration was performed for both positive and negative ionization modes using a standard calibration mix (Thermo Fisher Scientific). Lock-mass correction was applied throughout the run to improve mass accuracy. Quality control (QC) samples, prepared by pooling equal volumes from all experimental samples, were injected periodically to assess instrument stability and analytical reproducibility.

##### Data Processing & Analysis

TraceFinder (version 5.1, ThermoFisher Scientific) or Xcalibur™ (ThermoFisher Scientific) were used for LC-MS data analysis. Compound identification across both platforms was based on accurate mass measurements (± 5 ppm), retention time (± 0.5 min), and matching isotope patterns to those of an in-house standard mix containing relevant metabolites.

TraceFinder was used for method development, data processing, semi-automated peak detection, and report generation, providing quantified peak areas. A compound database containing accurate masses was employed for method development. Retention times were verified using standard mixes containing relevant compounds. Peak areas were normalised against the internal standard peak area and either total protein content (µg), dry weight (mg), or volume (µL).

For fractional enrichment quantification and natural isotope abundance correction, a custom script (Python version 3.12) was used^35^. Fractional enrichment is presented as the percent molecule labelled.

#### Statistics and Reproducibility

Statistical analysis was performed using ANOVA (one-way and two-way) or two-tailed Student’s unpaired or paired t-test, as indicated in the figure legends. Data distribution was assessed for normality using Q-Q plots. For repeated measures ANOVA, as in proliferation assays and tumour growth measurements, sphericity was not assumed, and a Greenhouse-Geisser correction was applied. All post-hoc analyses were corrected for multiple comparisons using the relevant method (Tukey, Sidak, and Dunnett). A *p*-value ≤ 0.05 was considered statistically significant, and the exact *p*-values for significant differences are indicated within the figures. The number of samples, biologically independent experiments, and statistical tests used are detailed in the corresponding figure legends. No statistical methods were used to predetermine sample sizes, however, these were in line with previously published sample sizes^1,29,36^. All data visualization and statistical analyses were conducted using GraphPad Prism (version 10.1.1).

For heatmaps of metabolite abundance, where metabolites of varying abundances were displayed together, min-max scaling was applied. This method scales values between 0% and 100%, where 0% was set to represent the absence of a metabolite, and 100% represented its highest abundance across all samples.

Where possible, mice from the same litter were randomized into groups before diet allocation and tumour cell injection. Data were excluded only in cases of technical failure or if identified as a statistically significant outlier (q ≤ 0.05; ROUT (Robust Regression and Outlier Removal) test). In some instances, mammary tumours within a cohort developed ulcerations early, requiring euthanasia based on experimental humane endpoints. These mice were excluded from downstream analysis.

## Resource availability

Data from bulk RNA sequencing of mammary gland tumours and mammary glands can be found in GEO number GSE295796.

All unique/stable reagents generated in this study are available from the lead contact with a completed materials transfer agreement.

## Acknowledgements

This work was supported by the Francis Crick Institute, which receives core funding from Cancer Research UK, the UK Medical Research Council, and the Welcome Trust (FC001223). To facilitate open access, the author has applied a CC BY public copyright license to any Author Accepted Manuscript version arising from this submission. We extend our sincere thanks to the animal technicians at the Francis Crick Biological Research Facility for their dedication and expertise in supporting this work.

## Author Contributions

E.G. and M.Y planned the experiments and wrote the manuscript. E.G. performed and analyzed experiments. P.C.D ran the LC-MS and NMR and provided intellectual input. A.M. and J.T.C. helped with experiments and provided intellectual input. M.K. contributed to GSE295796 and the RNA-seq analysis. S.S. provided essential animal husbandry support, including tumour measurements, mouse monitoring, and diet management. J.I.M. supervised the LC-MS and provided intellectual input.

## Supplemental Information

Document S1 containing Figures S1-S9.

