## Supplemental Figures S1-S9 for "Histidine exchange sustains LAT1 activity and proliferation in glutamine-addicted breast cancers"

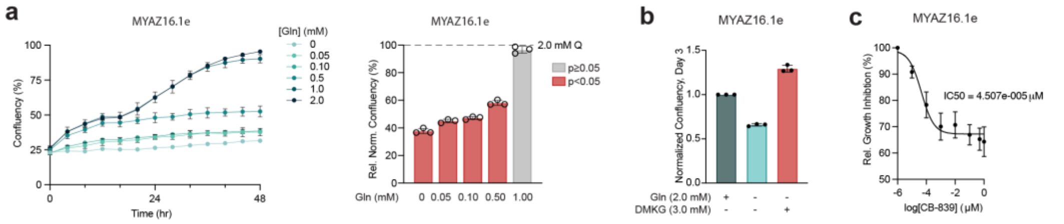


**Extended Fig. S1: Assessing Gln dependency in MYAZ16.1e MT cells.** (**a**) Confluency over time (left) of cells in 0-2.0 mM Gln and confluency normalized to 2.0 mM Gln at 48 h (right) (n = 3). Mean ± s.d.; *p*-values from two-way ANOVA (Šídák's correction). (**b**) Confluency normalized to 2.0 mM Gln at 72 h (n = 1). Mean ± s.d. (**c**) $IC_{50}$ curve for CB-839 growth inhibition (n = 1), from non-linear fit of log[CB-839] vs. normalized response.

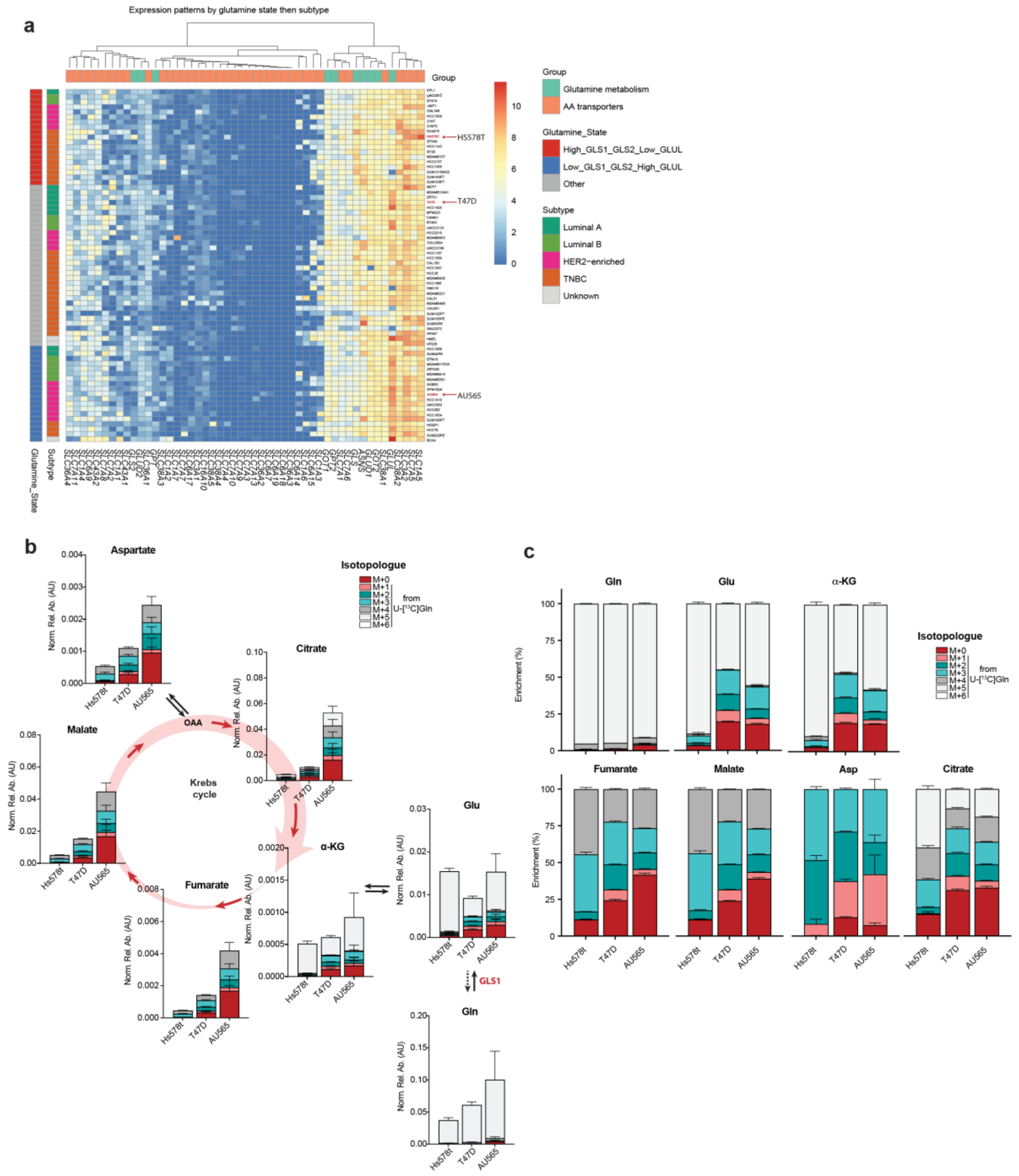


**Extended Fig. S2: Assessing Gln metabolism in human breast cancer cell lines.** (**a**) Heatmap of the expression of Slc and glutamine metabolism genes in breast cancer cells lines from DepMap (https://depmap.org/portal/). (**b**,**c**) Total abundance (**b**) and 13C isotopologue fractional enrichment (**c**) of indicated metabolites measured by LC-MS in cells cultures in the presence of 2mM [$U^{13}C$]Gln (24h, n = 3). Data are mean ± s.d.; *p*-values from two-way ANOVA (Šídák’s correction).

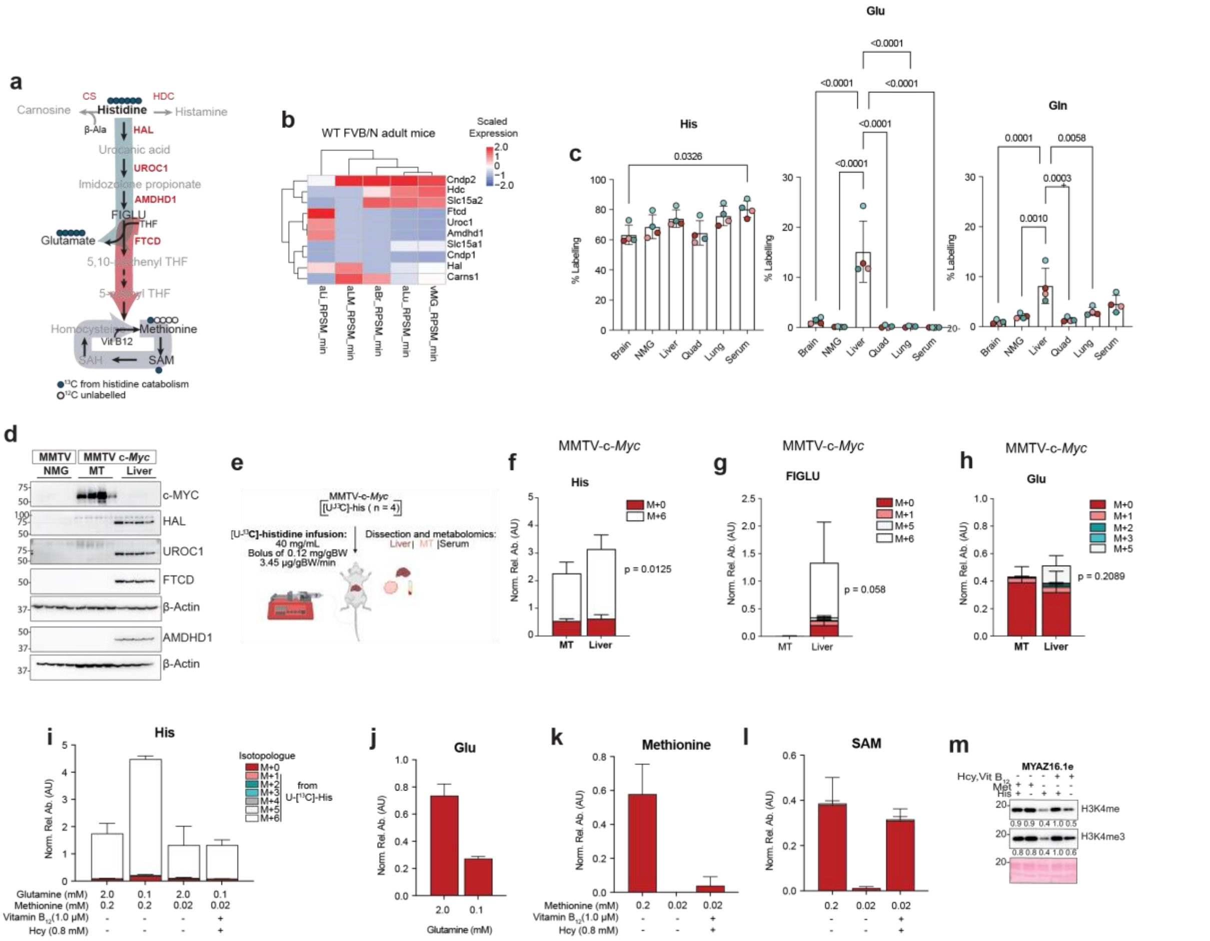


**Extended Fig. S3: Assessing His catabolism in tissues and MYC-driven tumour cells.** (**a**) Schematic of His catabolism. (**b**) Heatmap of His metabolic enzyme Z-score normalized expression from GSE75957. *aLi*: adult liver; *aLM*: adult lung; *aBr*: adult brain; *vMG*: virgin mammary gland. (**c**) LC-MS of $^{13}$C enrichment in WT mice after a [U$^{13}$C]His infusion (3 h). Mean ± s.d.; *p*-values from two-way ANOVA (Šídák's correction). (**c**) LC-MS of $^{13}$C enrichment in WT mice after a [U$^{13}$C]His infusion (3 h). Mean ± s.d.; *p*-values from two-way ANOVA (Šídák's correction). (**d**) WB of His catabolic enzymes in NMG (WT controls, n = 4), MMTV-c-*Myc* tumours (n = 4), and liver tissue. (**e**) [U$^{13}$C]His infusion strategy in MMTV-c-*Myc* mice. (**f-h**) LC-MS of metabolite abundance in tumours and matched liver samples from MMTV-c-*Myc* mice (n = 4). Mean ± s.d.; *p*-values from two-way ANOVA (Šídák's correction). (**i-l**) LC-MS of metabolite abundance in MYC-driven tumour cells (n = 3) cultured with 0.2 mM [U$^{13}$C]His (24 h). (**m**) WB of histone extracts from MYC-driven tumour cells (n = 3) under indicated conditions, protein levels were normalized to Ponceau staining and expressed relative to the maximum value.

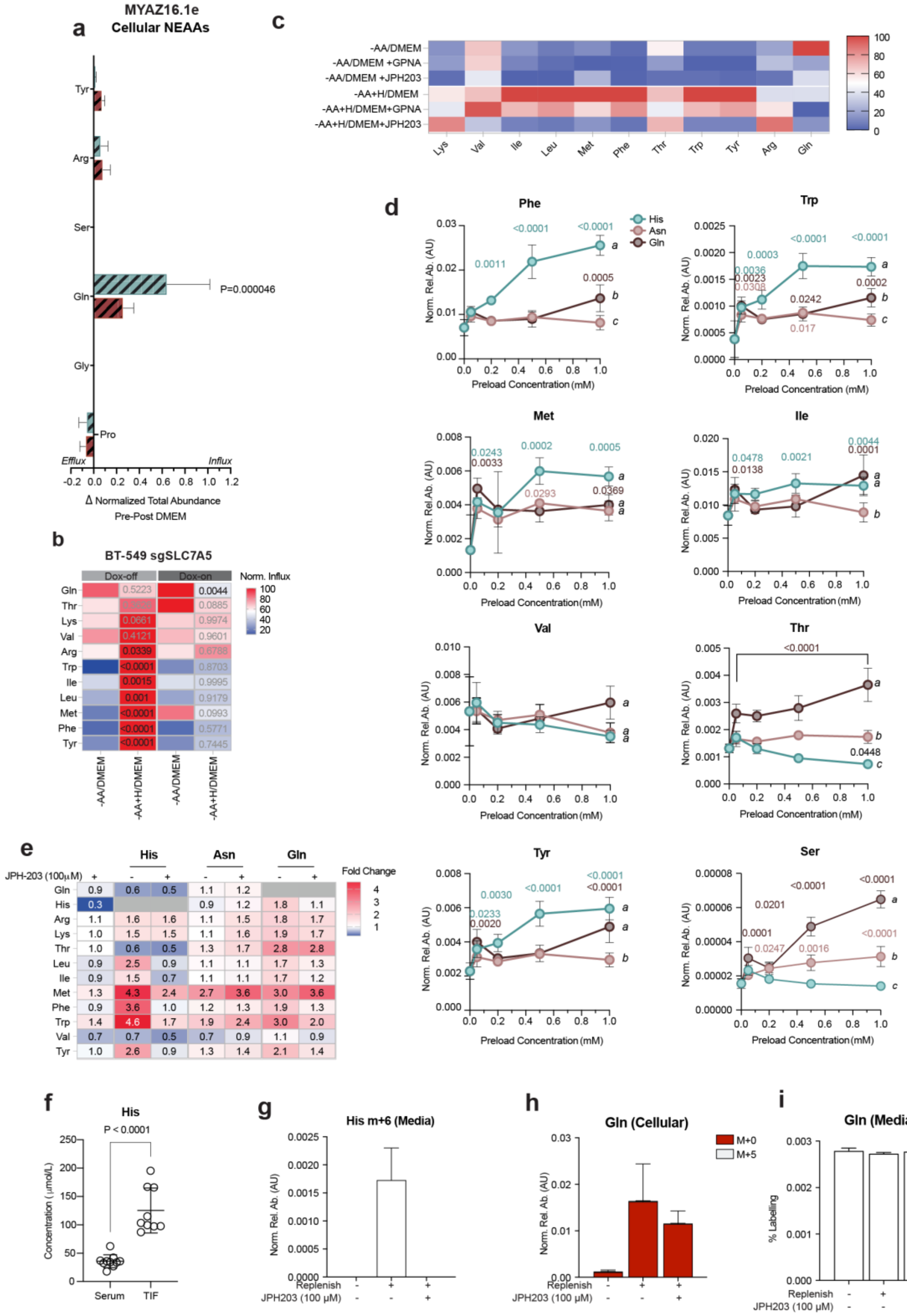


**Extended Fig. S4: LC-MS analysis of His-preloaded cells.** (**a**) Intracellular NEAA flux in cells preloaded with 1.6 mM His (48 h) and replenished with DMEM for (2.5 min, n = 3). Mean ± s.d.; *p*-values from two-way ANOVA (Šídák's correction). (**b**) Heatmap of amino acid influx, normalized to the maximum per amino acid, in His-preloaded BT-549 sgSLC7A5 cells (48 h, 1.6 mM), replenished with DMEM (2.5 min; n = 3). (**c**) Heatmap of intracellular amino acid abundance in cells preloaded with 1.6 mM His and replenished ± inhibitors (2.5 min), normalized to the maximum per amino acid. (**d**) Amino acid uptake in cells preloaded with His, Asn, or Gln (48 h), then replenished with DMEM (2.5 min, n = 3). Different letters indicate significant differences. (**e**) Heatmap of intracellular amino acid abundance in starved or preloaded cells (1.0 mM His, Asn, or Gln), replenished with DMEM ± JPH203 (2.5 min, n = 3). Colour scale represents mean fold change vs. starved cells, with values displayed. All statistical tests are mean ± s.d.; *p*-values from two-way ANOVA (Šídák's correction). (**f**) Absolute His concentrations in serums (n = 10) and TIF (n = 9) from mice bearing MYAZ16.1e orthotopic tumours. Mean ± s.d., paired two-tailed t-test. (**g-i**) Cells preloaded with 0.05 mM [$U^{13}C$]His and 0.3 mM [$U^{13}C$]Gln (48h), replenished ± JPH203 for 2.5 min (n = 3): (**g**) Relative m+6 His abundance in media, (**h**) Relative Gln abundance in cells, (**i**) Percent labelled Gln in media.

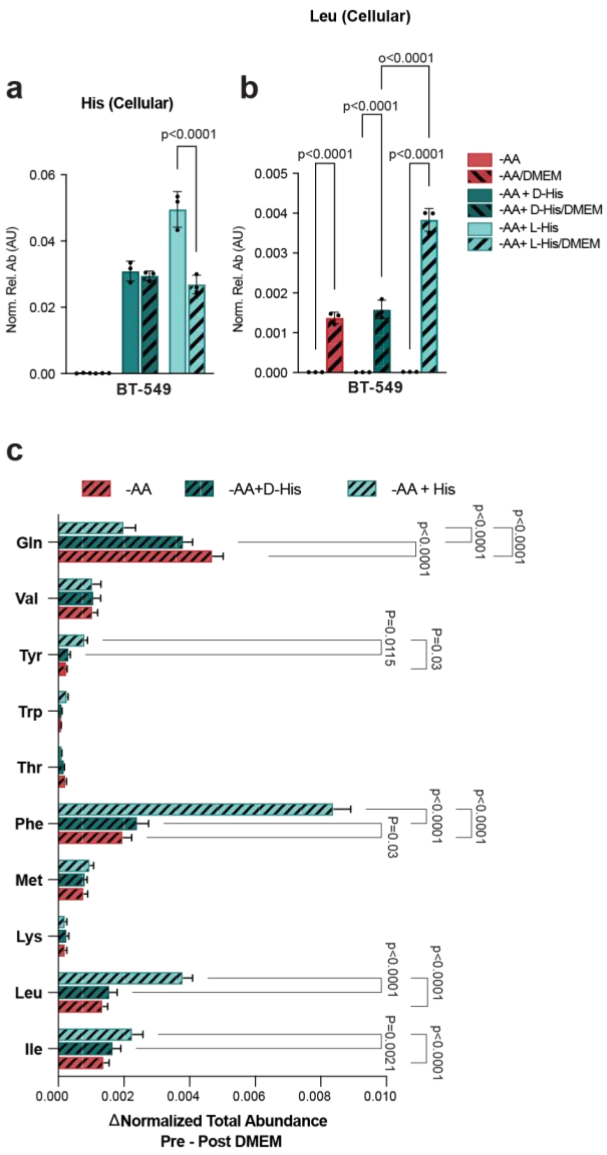


**Extended Fig. S5. LC-MS analysis of D-His preloading assay.** Cells starved of amino acids (-AA) ± 1.6 mM D-His preload (-AA+D-His) for 48 h, then replenished with DMEM (2.5 min, n = 3). (**a,b**) Relative total intracellular abundance of (**a**) His and (**b**) Leu. (**c**) Relative influx of amino acids, calculated as the difference pre and post DMEM relative total abundances. Mean ± s.d.; *p*-values from two-way ANOVA (Tukey's correction).

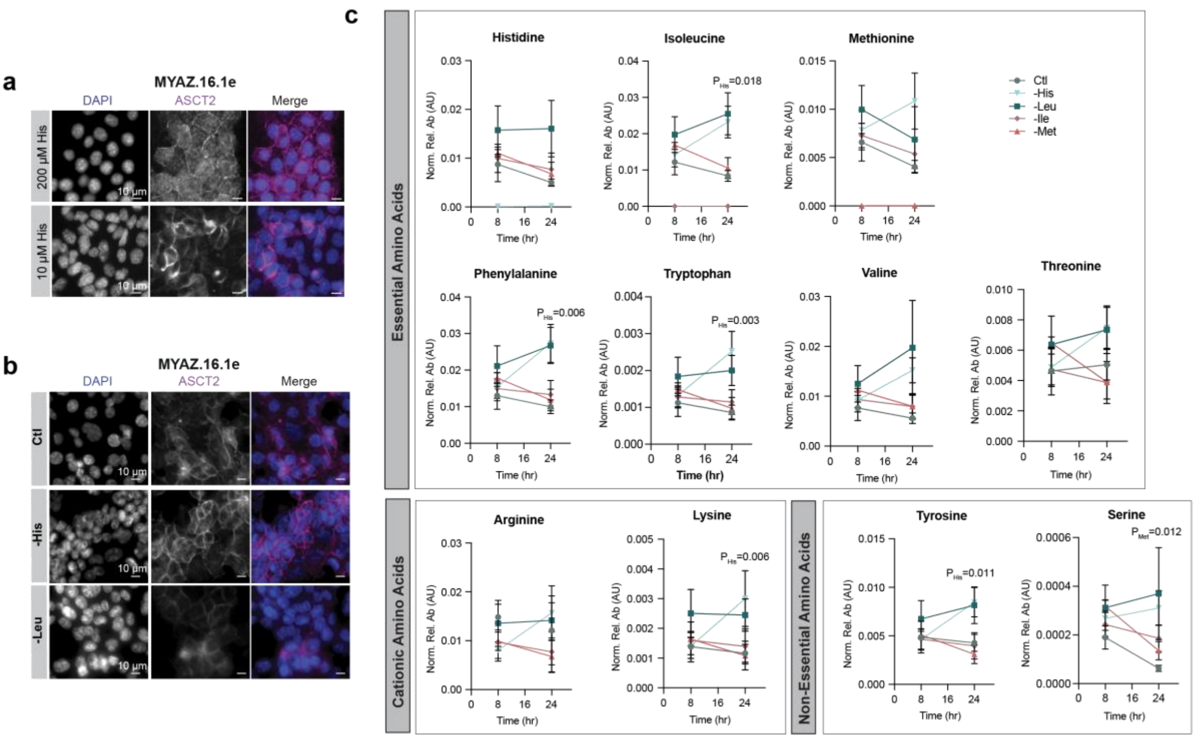


**Extended Fig. S6: Effects of His depletion.** (**a**,**b**) Immunofluorescence images of ASCT2 localisation in cells cultured with (**a**) 200 µM or 10 µM His (16 h) and (**b**) complete (Ctl), His-, or Leu-free DMEM (16 h). (**c**) LC-MS of intracellular amino acids in cells cultured in Ctl, or His-, Leu-, Ile-, or Met-free DMEM (n = 3), comparing 8 h to 24 h. Mean ± s.d.; *p*-values from two-way ANOVA (Šídák's correction).

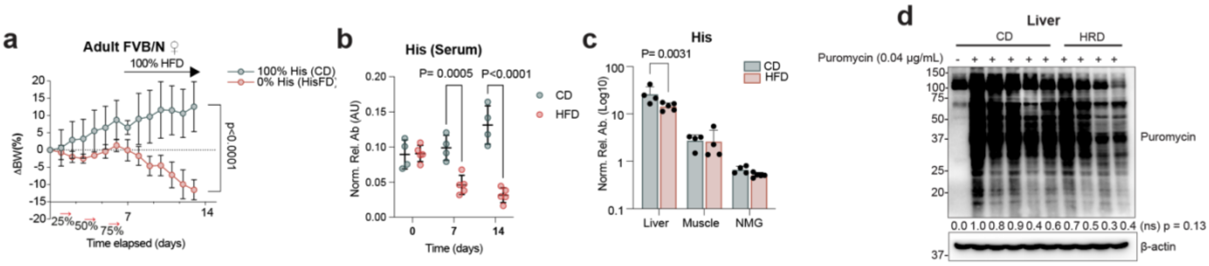

**Extended Fig. S7: Effects of HisFD and 20%HRD on healthy mice.** (**a-c**) Wildtype FVB/N mice were on a CD (n = 4) or HisFD (n = 5). (**a**) Body weight change over time. (**b,c**) LC-MS of His abundance in (**b**) serum over time and (**c**) tissues. *P*-values from two-way ANOVA (Šídák's correction). **d**) WB of liver tissue from a subset of CD (n =5) and HRD (n=4) mice injected with puromycin (Puro), and an uninjected CD-mouse. Protein levels were normalized to β-actin and expressed relative to the maximum. *P*-values from unpaired two-tailed t-test. All data mean ± s.d.

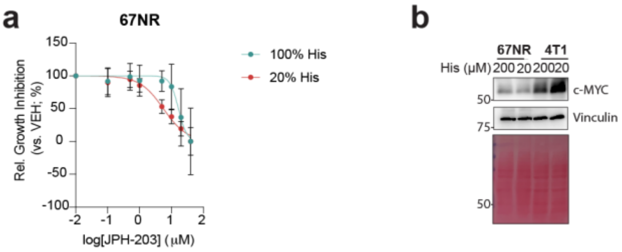


**Extended Fig. S8: His limitation affects MYC levels and the sensitivity to JPH203.** (**a**) IC50 curve for JPH-203 in 67NR cells cultured in high or low His for 72 hours (n = 3). Data are mean ± s.d.; *p*-values from two-way ANOVA with Šídák's correction. (**b**) WB of MYC expression in 67NR and 4T1 cells cultured in high and low His (n = 1).

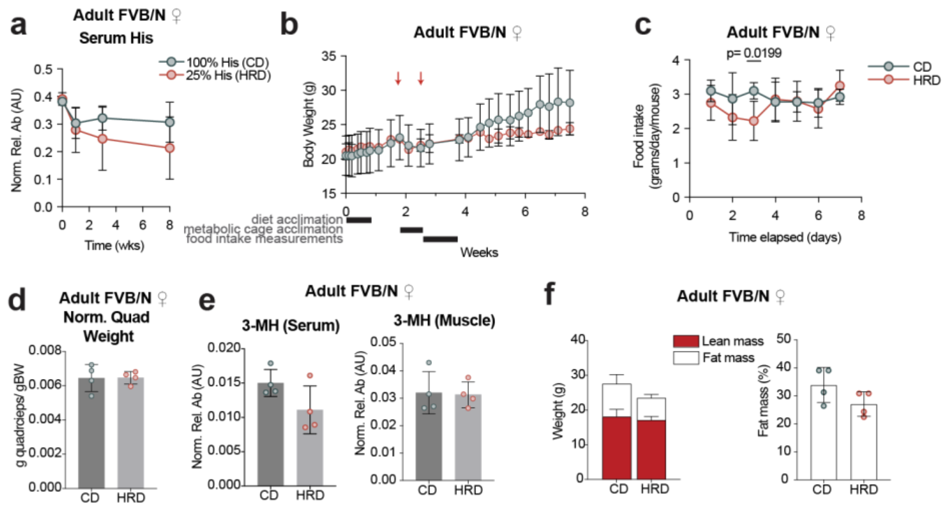


**Extended Fig. S9: Effects of 25%HRD on healthy mice.** (**a-f**) Wild-type FVB/N mice on a 25%HRD (n = 4) or CD (n = 4). (**a**) LC-MS of serum His levels over time. (**b**) Body weight over time, arrows indicate individual housing in metabolic cages, which caused minor weight loss. *P*-values from two-way ANOVA (Šídák's correction). (**c**) Food intake. *P*-values from two-way ANOVA (Šídák's correction). (**d**) Quadriceps muscle weight normalized to total body weight. *P*-values from unpaired two-tailed t-test. (**e**) LC-MS analysis of 3-MH in serum (left) and quadricep muscle (right). *P*-values from unpaired two-tailed t-test. (**f**) Body composition: proportion of lean and fat mass (left) and change in fat mass percentage (right). *P*-values from unpaired two-tailed t-test. All data mean ± s.d.
